# Reconstructing the regulatory programs underlying the phenotypic plasticity of neural cancers

**DOI:** 10.1101/2023.03.10.532041

**Authors:** Ida Larsson, Felix Held, Gergana Popova, Alper Koc, Rebecka Jörnsten, Sven Nelander

**Author notes:** Indicates equal contribution (Ida Larsson, Felix Held).

## Abstract

Nervous system cancers contain a large spectrum of transcriptional cell states, reflecting processes active during normal development, injury response and growth. However, we lack a good understanding of these states’ regulation and pharmacological importance. Here, we describe the integrated reconstruction of such cellular regulatory programs and their therapeutic targets from extensive collections of single-cell RNA sequencing data (scRNA-seq) from both tumors and developing tissues. Our method, termed single-cell Regulatory-driven Clustering (*scRegClust*), predicts essential kinases and transcription factors in little computational time thanks to a new efficient optimization strategy. Using this method, we analyze scRNA-seq data from both adult and childhood brain cancers to identify transcription factors and kinases that regulate distinct tumor cell states. In adult glioblastoma, our model predicts that blocking the activity of *PDGFRA*, *DDR1*, *ERBB3* or *SOX6*, or increasing *YBX1* -activity, would potentiate temozolomide treatment. We further perform an integrative study of scRNA-seq data from both cancer and the developing brain to uncover the regulation of emerging meta-modules. We find a meta-module regulated by the transcription factors *SPI1* and *IRF8* and link it to an immune-mediated mesenchymal-like state. Our algorithm is available as an easy-to-use R package and companion visualization tool that help uncover the regulatory programs underlying cell plasticity in cancer and other diseases.

## Introduction

Nervous system cancers in adults and children share a common trait: tumor cells exist in multiple transcriptional states, which partly resemble the diversification of cells during normal embryonic development. In adult glioblastoma (GBM), cells resemble neural progenitor cells (NPCs), oligodendrocyte progenitor cells (OPCs) or astrocytes (ACs) (Neftel *et al*, 2019). In addition to these states, a substantial fraction of cells display a mesenchymal (MES)/injury response-like profile that have been linked to monocyte populations (Richards *et al*, 2021; Wang *et al*, 2017; Gangoso *et al*, 2021; Schmitt *et al*, 2021; Hara *et al*, 2021). In childhood cancers such as diffuse midline glioma, tumor cells move from a proliferative, OPC-like state to more differentiated states resembling either AC- or OC-like lineages (Filbin *et al*, 2018), and in medulloblastoma (MB) the four main subgroups differ in the proportion of undifferentiated vs differentiated tumor cells and their lineage resemblance (Hovestadt *et al*, 2019).

There are strong reasons to believe that the cell states found in a tumor have implications for disease progression. For instance, invasive GBM cells mimic the migration mechanisms of neuronal cells (Cuddapah *et al*, 2014; Venkataramani *et al*, 2022), whereas tumor initiation is mediated by quiescent stem-like cells (Xie *et al*, 2022). Further, it’s been reported that recurrent therapy-resistant GBM tumors display a higher fraction of MES cells (Bhat *et al*, 2013), possibly due to a phenotypic shift in the cells from non-MES to MES (Wang *et al*, 2022). However, despite their role in disease recurrence, progression, and drug resistance, we have limited knowledge about the mechanisms by which cell states are regulated. In particular, to understand the transcriptional regulation it is essential to identify sets of transcription factors (TFs) whose activity have a direct impact on a specific state phenotype. Furthermore, to identify drug targeting opportunities, it is important to identify sets of druggable proteins, particularly kinases, that can be linked to states.

Amounting repositories of scRNA-seq data presents new opportunities to uncover such regulation with high precision. However, existing data analysis methods are not developed with this goal in mind. The frequently used clustering methods do not contain regulatory predictions; they are only concerned with grouping cells or genes. Previously, a standard method to identify regulators of gene signatures has been to first estimate a gene regulatory network (GRN), followed by post-processing to identify regulators linked to signature genes. This approach can, for instance, predict regulators of mesenchymal transformation from bulk transcriptional (Carro *et al*, 2010) and bulk multi-omic (Kling *et al*, 2016) data, but transferring it to the scRNA-seq setting has proven unsatisfactory as individual links are hard to reproduce (Pratapa *et al*, 2020; Chen and Mar, 2018). Adding additional layers of data, primarily ATACseq or CHIPseq measurements (Aibar *et al*, 2017; Kamimoto *et al*, 2020; Kamal *et al*, 2022), is one way of alleviating the problem but comes with its own limitations. Existing methods are not fast enough to process millions of cells, and restricting the model to only TFs makes it less amenable to modeling the impact of pharmacologically more relevant gene classes, such as kinases. Thus, the development of fast and accurate methods to reconstruct gene regulatory programs from scRNA-seq data will be important, if not essential.

Here, we describe a new method for fast construction of regulatory programs that is well-suited for large data sets and for the detection of critical regulators of cell states. Applied to scRNA-seq data, it jointly detects modules (sets, clusters) of co-expressed genes and groups of regulators, such as transcription factors and kinases, linked to each module. Compared to methods of similar scope it is both faster and more flexible. As proof-of-principle, we show the applicability of our algorithm through three use cases. First, we apply it to a data set from peripheral blood mononuclear cells (PBMC) and benchmark it against the most comparable algorithm we can find, SCENIC+ (González-Blas *et al* (2022)). Second, we apply it to scRNA-seq data previously generated by our group (Larsson *et al*, 2021) to predict regulatory interventions to potentiate temozolomide (TMZ) treatment in GBM. Third, we integrate 13 data sets from the developing brain and cancers of the nervous system to investigate the regulatory landscape of neuro-oncology. We find a meta-module regulated by the transcription factors *SPI1* and *IRF8* which show strong resemblance with the previously described immune cell-induced MES-like state. The algorithm we developed, *scRegClust*, is available as an easy-to-use R-package.

## Results

### A regulatory-driven clustering method

We present a new approach to detect regulatory programs from scRNA-seq data alone, based on a mathematical framework that directly models the interaction between regulators and responding target gene sets (Figure 1A). Our model assumes that genes belong to either of two categories: tentative regulators and tentative targets. Given a volume of data **Z**, organized as cells × genes, we split the data in two parts, a regulatory part **Z***_r_*, and a target part **Z***_t_*. These parts are further split randomly into two sets of training and assessment cells. We then perform a clustering task by identifying groups of genes in **Z***_t_* that correspond to sets of co-regulated genes, and for each such set, we perform a regulatory program re-construction task by identifying a small number of genes in **Z***_r_* that are the likely regulators. These steps are repeated until configurations stabilize. To fit regulatory programs from data computationally, we developed an alternating two-step scheme iterating between determining the most predictive regulators (regulatory program re-construction) for each of a pre-specified number *K* of modules and using these optimal regulators to allocate target genes into *K* modules (clustering). Step 1 is approached using cooperative-Lasso (coop-Lasso, Chiquet *et al*, 2012) on the training set. In this step, we search for a sparse set of regulators that can be linked to most genes in a module, each regulator with the same sign (positive vs negative). A sparsity penalty parameter controls the number of regulators assigned to each module. In section “Regulatory-driven clustering improves accuracy” we provide details on how to choose this penalty parameter. To solve this optimization problem efficiently, we propose an adaptation of over-relaxed Alternating Direction Method of Multipliers (ADMM, Boyd *et al*, 2011; Xu *et al*, 2017). Once regulators have been assigned to modules, we move on to Step 2 where the task is to refine the target gene sets. We approach this by re-estimating coefficients per module using sign-constrained non-negative least squares (NNLS, Meinshausen, 2013) on the training set and an observation-based allocation scheme using the assessment set. Target genes whose predictive *R*^2^ is below a threshold across all modules are marked as noise and placed in a rag-bag cluster. This ensures that outliers do not overly distort the clustering result. There are several optional inputs that can guide the algorithm. Of note is the possibility to provide prior information on the relationship between genes in the form of a gene × gene matrix where a non-zero entry indicates that there is a known link between the genes. A known link can for example be that both genes appear in the same biological process. Algorithmic details can be found in “Methods”. The resulting regulatory programs are visualized using either a coupling matrix view, where predicted regulators are columns and target gene modules are rows, or a module network view (Figure 1A). In addition, the algorithm returns goodness-of-fit measures which can be used to guide the selection of optimization parameters (Figure 2A, Figure S6).

**Figure 1.**
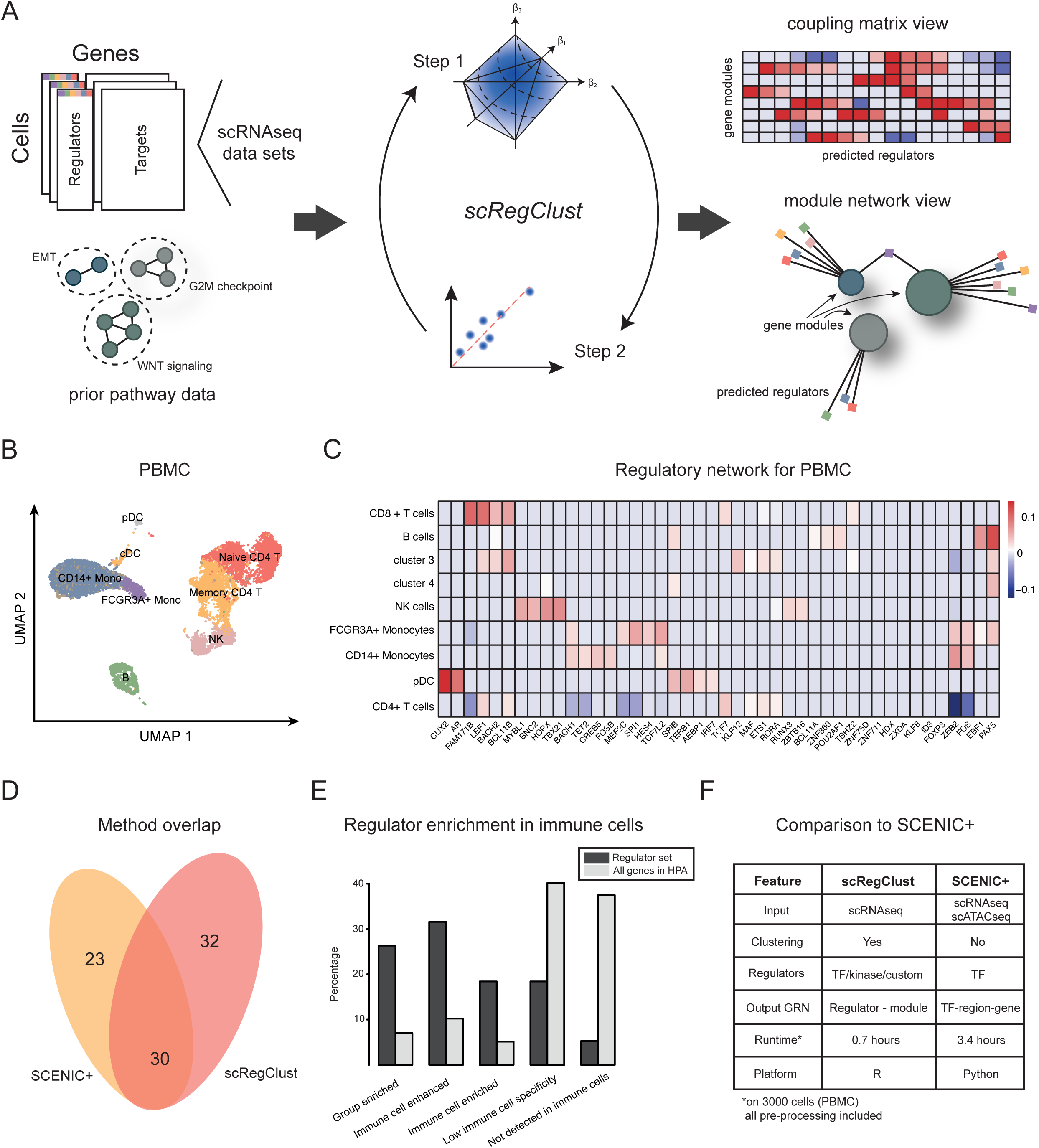
A regulatory-driven clustering method. **A** A schematic overview of the algorithm pipeline from gene expression data to regulatory networks. **B** UMAP embedding of the PBMC cells colored according to immune cell type assignment. **C** The regulatory table inferred by *scRegClust*. Rows correspond to immune cell types and columns to regulators (TFs). The scale goes from high positive regulation (red) to high negative regulation (blue), as indicated by the key. **D** Venn diagram showing the overlap in identified regulators between SCENIC+ and *scRegClust*. **E** Regulator enrichment in the Human Protein Atlas compared to a baseline of all genes in the atlas. **F** Comparison between *scRegClust* and SCENIC+.

**Figure 2.**
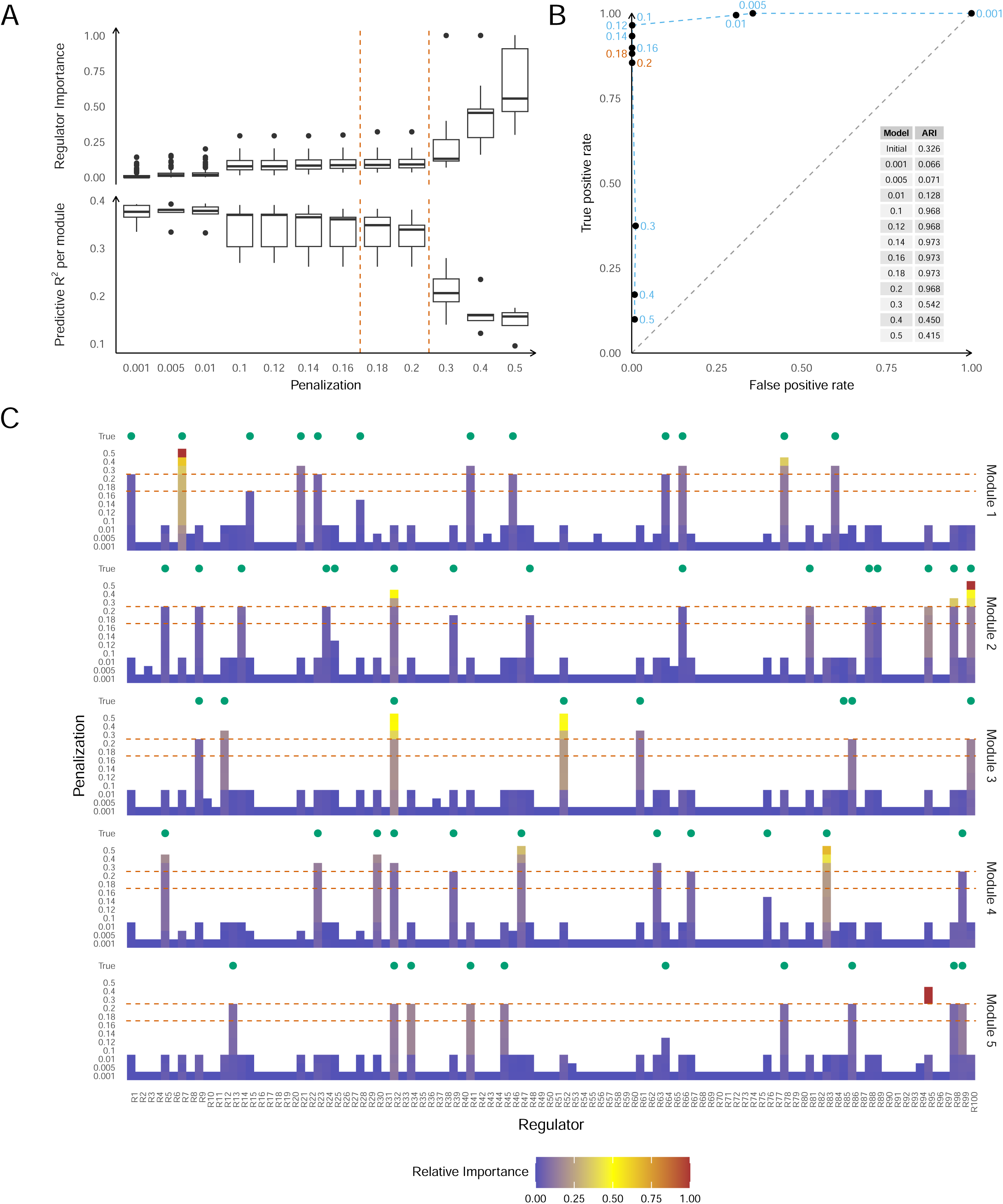
Method validation and analysis tools. Results from a representative run of *scRegClust* on the data from simulation setup A (see the Supplementary Material for details on data generation). **A** Boxplots of predictive R^2^ per module and importance per regulator shown across a progression of penalization parameters. Dashed lines indicate a region of solutions that demonstrates our selection rule. **B** Average true positive rate (TPR) shown against average false positive rate (FPR) in a ROC curve, illustrating the quality of groundtruth regulator identification. TPR and FPR are computed per target gene and then averaged. The corresponding penalization parameter is shown with each average. A table of ajusted Rand indices between the estimated clustering and the true clustering is shown as an inset. **C** Relative importance of the regulators within each module as estimated by *scRegClust* within each penalization level. Each plot represents one of the five estimated modules and each line shows the result for a penalization parameter. Colored tiles indicate that the regulator was included in the final model for the respective module. Green dots above the regulators for each module indicate regulators used during simulation of the data. The dashed lines correspond to the dashed lines in part A. They illustrate which regulators would be selected when the penalization parameter was chosen by our selection rule.

### Comparison with SCENIC+ shows good agreement between methods

We benchmarked our entire workflow on a publically available data set from peripheral blood mononuclear cells (PBMC). Starting from a matrix of raw gene counts, the data was normalized and formatted (see details in “Methods”). As previously mentioned, potential regulators can be all TFs, all kinases, or any custom assignment of interest. For a direct comparison with the newly released SCENIC+ (González-Blas *et al*, 2022) (combining scRNA-seq and scATAC-seq), we used all TFs as potential regulators. We did not provide the algorithm with any prior information. Once the algorithm converged, it had clustered the PBMC data into nine gene modules and 62 active TFs interacting with one or several of these gene modules.

From previous work on the PBMC data, we know that the cells can be divided into 8 clusters representing various types of immune cells (Figure 1B, Figure S1A). To functionally annotate the gene modules defined by *scRegClust*, we scored these against the gene signatures of the immune cell types (Figure S1B-C). Our algorithm can successfully distinguish modules upregulated in CD8+ T cells, CD4+ T cells, B cells, NK cells, FCGR3A+ monocytes, CD14+ monocytes and plasmacytoid dendritic cells (pDCs). Among the strongest regulators we find many examples of known regulators, e.g. *PAX5* (B cells), *TBX21* (NK cells), *CUX2* (dendritic cells) and *LEF1* (T cells), to mention a few (Figure 1C). As a point of comparison, SCENIC+ identified 53 (activator) TFs, out of which 30 overlap with our identified regulators (*p* < 10^−5^, Fisher’s exact test; Figure 1D). To get a more detailed understanding of our output, we looked into the functional annotation for the predicted regulators with a particular focus on the 32 TFs that do not overlap with SCENIC+. For the regulators that had an annotation in the Human Protein Atlas (HPA, Uhlén *et al*, 2015), 76.3 % are classified as being immune cell enriched/enhanced compared to a baseline 22 % for all genes in the atlas. For the non-overlapping genes, 55 % are enhanced or enriched in immune cells and have an immune cell type annotation that agrees with the one predicted by *scRegClust* (Figure 1E, Supplementary Table 1).

Jointly, the comparison between *scRegClust* and SCENIC+ on PBMC data showed a good level of agreement, despite the fact that *scRegClust* uses scRNA-seq data alone. In contrast to SCENIC+, which relies on pre-defined cell types for each cell in the primary data, *scRegClust* performs clustering on target genes, which can then be functionally annotated to match cell types. As previously mentioned, SCENIC+ is restricted to identifying TFs as regulators, due to its reliance on scATAC-seq data, while *scRegClust* in principle can consider any category of genes as potential regulators. The speed of the two algorithms is also remarkably different, *scRegClust* being three times faster than SCENIC+ (Figure 1F).

### Regulatory-driven clustering improves accuracy

We also benchmarked our workflow on synthetic data to explore the effect of associating regulators with modules on clustering quality, the robustness of our modeling assumptions, and correct regulator recovery. In addition, we developed tools and strategies to guide the selection of penalty parameters and the correct number of modules. The quality of the clustering with respect to the groundtruth is measured using the adjusted Rand index (Hubert and Arabie, 1985). Two measures are introduced to evaluate the quality of the selected regulatory model associated with each module. Predictive *R*^2^ per module measures the predictive performance on the assessment set of the regulators associated with each module. Regulator importance measures the change in predictive *R*^2^ per module when a single regulator is omitted from the full regulator model for a module. In addition, we considered the true and false positive rates, presented as a ROC curve, of estimating the correct regulators associated with each target gene. See the “Methods” section for details on the computed metrics.

Synthetic data was generated to mimic characteristics of our in-house scRNA-seq data as well as characteristics of the multi-study comparison presented in the upcoming sections. We based our simulations on a modified scRNA-seq data set (see “Supplementary Methods”) and generated four different setups capturing different aspects of real data scenarios. Below, a summary of the most important findings is presented. Consult the Supplementary Material for details on simulations and further results.

If no initial clustering is provided, our algorithm is initialized by k-means++ (Arthur and Vassilvitskii, 2007) on the cross-correlation matrix of the target genes and regulators in the training set. When the penalization parameter was chosen appropriately, associating regulatory models with each module improved the clustering quality substantially (Figure 2B, adjusted Rand index increases from 0.33 to 0.97). *scRegClust* showed excellent regulator recovery properties in our simulations (see Figure 2B and C). For appropriate penalization, the correct regulators were selected and only few true regulators were disregarded. *scRegClust* performed well even when model assumptions were violated (see the Supplementary Material). Notably, *scRegClust* achieved a large improvement in clustering quality in a scenario with unequal module sizes. There, clustering on the cross-correlation matrix gave an essentially random result, whereas our algorithm managed to nearly correctly recover the simulated modules, when penalization was chosen appropriately (see Figure S5B, adjusted Rand index increases from 0.06 to 0.95). This shows that the integrated approach of structure modeling and clustering is superior to clustering on the correlation structure alone.

Predictive *R*^2^ per module and regulator importance can be used as guidance for the selection of the appropriate range of penalty parameters. As a general rule, at least 5 to 10 different penalization parameters should be tested. Increasing the penalty parameter reduces the number of active regulators associated with each module. This will typically reduce predictive *R*^2^ gradually until the point when penalization becomes too strong and important regulators are not associated with a module any longer. A similar but reversed trend can be observed for regulator importance. Due to many regulators being included in the models for low penalization, importance will typically be low at first. Increasing the penalty parameter and therefore including fewer regulators in the model will increase regulator importance until it becomes inflated due to the inclusion of too few regulators. As illustrated in Figure 2A, finding this change point from gradual to rapid decrease or increase, respectively, indicates a range of appropriate penalty parameters.

A common problem in clustering is the selection of the correct number of modules *K*. Contrary to methods like k-means, our algorithm can produce empty modules. This can be enforced more strongly by specifying a minimum non-empty cluster size. We find in our simulations that *scRegClust* tends to combine groundtruth modules with regulatory overlap when *K* is underspecified and tends to split groundtruth modules into smaller ones when *K* is overspecified. To provide further guidance on the selection of the number of modules, we introduce silhouette scores (see “Methods” section). These can be interpreted similarly to standard silhouette values (Rousseeuw, 1987) as a measure of how well a target gene is located within its own module compared to how well it would be located within the nearest module. High average silhouette scores indicate that target genes are well located within their module. Analysis of the average silhouette score should be combined with analysis of the predictive *R*^2^ per module (see Figure S6). A good clustering achieves high values in both scores. Our recommendation is to start with an initial guess for the number of modules *K*_0_ and to run *scRegClust* on a range of module counts above and below *K*_0_. Analysis of the silhouette scores and predictive *R*^2^ per module as illustrated in Figure S6 can give guidance on how to choose the optimal number of modules.

To summarize, through an extensive simulation study where several data simulation setups reflecting various real data scenarios were tested, we demonstrated that *scRegClust* performs excellent both in terms of improving clustering accuracy and recovering regulatory interactions.

### Targeting regulators of an OPC-like state potentiates temozolomide treatment

Encouraged by the results from the benchmarking studies, we continued by applying *scRegClust* to complex data sets from nervous system cancers to explore two key questions. First, we asked if the algorithm could predict interventions, such as drugs or gene perturbations, that would push GBM cells into a temozolomide-sensitive state. In subsequent passages, we leverage the power of our algorithm to integrate scRNA-seq data across nervous system cancers and the developing brain to identify meta-modules with shared modes of regulation and biological function.

One of the questions raised during our previous work on cell state transitions in GBM was how to successfully target an entire cell state, and not just individual target genes of these states. The question was prompted by our prediction that the minimal intervention needed to potentiate temozolomide (TMZ) treatment is to block transitions to what we call “state 5”, a state with an OPC-like, invasive profile (Larsson *et al*, 2021) (Figure 3A). To investigate this, we applied *scRegClust* to the previously mentioned scRNA-seq data from U3065MG cells (Xie *et al*, 2015) using either TFs or kinases as potential regulators (Figure 3B-C). The algorithm was run without giving prior information on cluster assignment for the genes, which resulted in *scRegClust* clustering the genes in 9 (TF) or 6 (kinase) modules, respectively. Important to note is that the six cell states are defined by clustering **cells** while *scRegClust* clusters **genes**, hence a discrepancy in module-state correspondence can be expected. The gene modules defined by our algorithm were related to the gene signatures derived from differential expression analysis of the six cell states (Figure 3A) and the overlap was scored using Jaccard index (Figure 3D).

**Figure 3.**
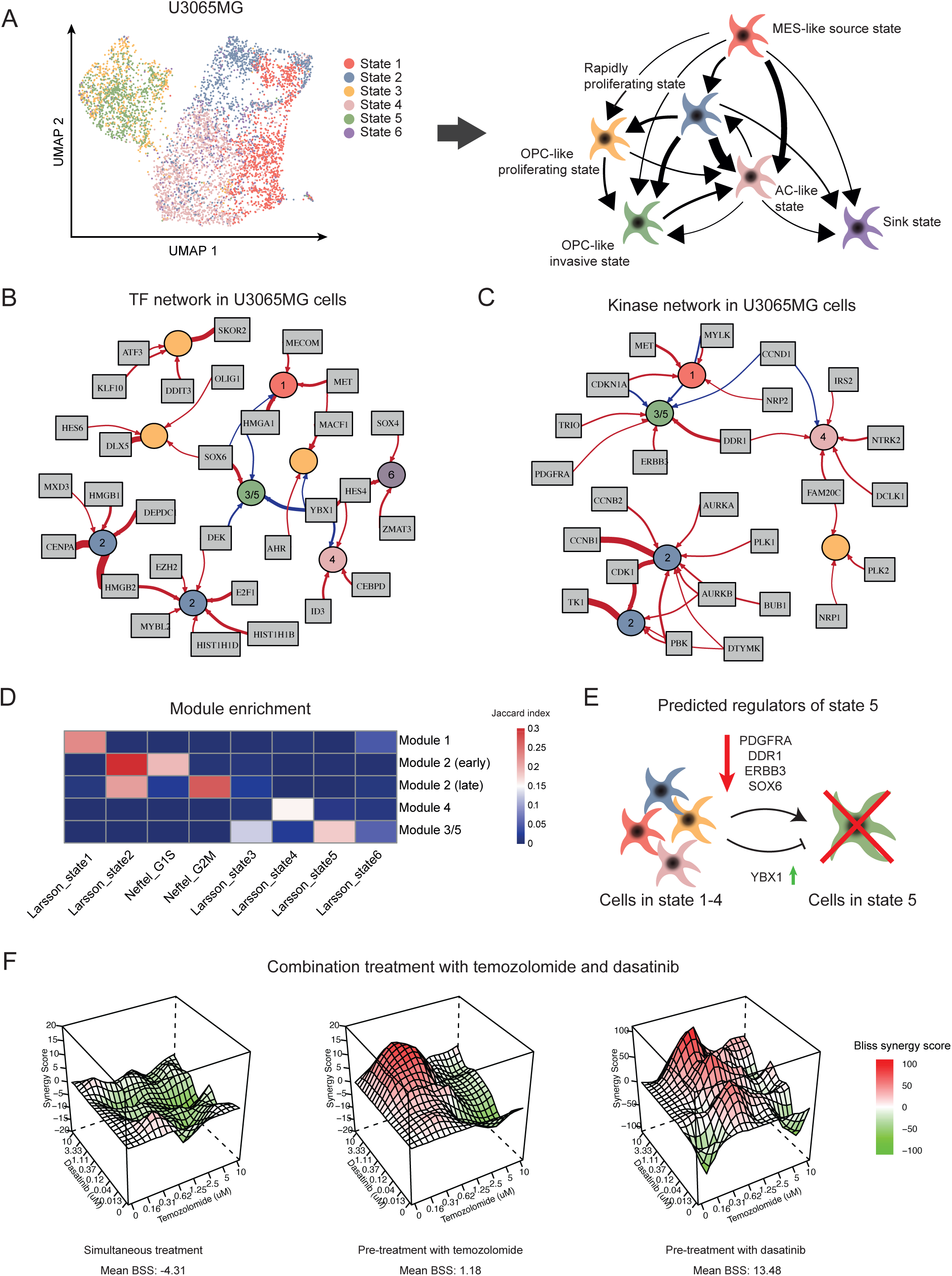
Regulatory interventions to potentiate TMZ treatment. **A** Left panel: UMAP embedding of the scRNA-seq data from (Larsson *et al*, 2021). Colors according to cell state assignment, as defined in the original paper. Right panel: The state transition network generated by the STAG model in the original paper, with state profile annotated according to gene set enrichment. **B-C** The regulatory networks generated by *scRegClust* for scRNA-seq data from (Larsson *et al*, 2021), in transcription factor (B) and kinase (C) mode. **D** Module enrichment against state signatures from the original paper, as well as gene signatures of cell cycle phases. Enrichment is measured using Jaccard Index. **E** Schematic of the predicted regulation of state 5. *PDGFRA*, *DDR1*, *ERBB3* and *SOX6* positively regulated state 5 and should be knocked down to block transitions to state 5. *YBX1* negatively regulated transitions to state 5 and should be overexpressed to block transitions to state 5. **F** The synergy landscape of temozolomide and dasatinib for (i) simultaneous treatment only, (ii) pre-treatment with temozolomide and (iii) pre-treatment with dasatinib.

We found that *scRegClust* defines two modules that correspond to state 2, a highly proliferative progenitor state. Looking at the enrichment of the gene modules against a collection of neurooncology-related gene sets (Larsson *et al*, 2021) (Figure 3D), it is evident that both modules are related to proliferation and the cell cycle and that they diverge on cell cycle phase - where one module represents the earlier phases of cell cycle (G1, G1S, S) while the second module represents the later phases (S, G2M, M). Several regulators identified for these modules agree with the module profiles, e.g. *E2F1* and *TK1* regulating the earlier phases of cell cycle (Martínez-Balbás *et al* (2000); Chang *et al* (1998)) while *AURKA* and *CENPA* are mostly active during G2M/M-phase (Marumoto *et al* (2003); Zeitlin *et al* (2001)). Another interesting observations is the astrocyte-like state 4, which is predicted to be positively regulated by *ID3*, known to induce astrocyte differentiation through the BMP pathway signaling (Bohrer *et al* (2015)), and *NTRK2*, involved in astrocyte maturation through BDNP signaling.

Addressing our initial question on how to suppress state 5 to potentiate TMZ treatment we focus on the regulation of the modules corresponding to this state. The top kinase regulator is *PDGFRA*, consistent with state 5 having an OPC-like profile as *PDGFRA* is both a known regulator of OPCs in normal development (Ellison and de Vellis, 1994) and frequently amplified in the OPC-like GBM state (Neftel *et al*, 2019). Other strong kinase regulators are *DDR1*, a receptor tyrosine kinase expressed in oligodendrocytes in the developing brain (Vilella *et al*, 2019), and *ERBB3*, also implicated in the oligodendrocyte lineage development and the OPC-like state in GBM (Wang *et al*, 2020). The top TF regulators are *SOX6* (positive regulation) and *YBX1* (negative regulation). *SOX6* belongs to the group D family of SOX TFs, which have been shown to regulate several stages of oligodendrocyte development. The combined predictions from Larsson *et al* (2021) and *scRegClust* suggest that blocking the activity of *PDGFRA*, *DDR1*, *ERBB3* or *SOX6*, or increasing *YBX1* -activity, would potentiate TMZ treatment (Figure 3E). We tested these predictions experimentally. First, we combined TMZ with the tyrosine-kinase inhibitor dasatinib, which had an inhibition score of > 0.85 in the STITCH database for *PDGFRA* and *ERBB3*, and an interaction score of 0.889 with *DDR1*. In the U3065MG cell line, simultaneous treatment with TMZ and dasatinib showed no synergistic effect, neither did pre-treatment with TMZ followed by simultaneous treatment with both drugs. A synergistic effect was, however achieved when pre-treating cells with dasatinib for 72 hours, followed by combined treatment with both drugs (BSS > 10) (Figure 3F). Relating this to previous predictions, the results imply that pre-treatment with dasatinib depletes the state 5 population, while subsequent combined treatment inhibits cells from escaping to state 5 and therefore leaves the tumor population vulnerable to TMZ. These conclusions would need to be corroborated by mechanistic studies.

Building on these results, we broadened our investigation’s scope to explore regulators of cell plasticity in nervous system cancers. To that end, we included more GBM single-cell data sets, as well as data from other cancers of the nervous system and the developing/healthy brain.

### The regulatory landscape of neuro-oncology

We used *scRegClust* to integrate the regulatory programs from the developing brain (Couturier *et al*, 2020; Eze *et al*, 2021; Hamed *et al*, 2022; Luo *et al*, 2021; La Manno *et al*, 2021; Weng *et al*, 2019), normal adrenal gland (Kildisiute *et al*, 2021), GBM (Couturier *et al*, 2020; Darmanis *et al*, 2017; LeBlanc *et al*, 2022; Neftel *et al*, 2019; Wang *et al*, 2019), MB (Hovestadt *et al*, 2019; Luo *et al*, 2021; Ocasio *et al*, 2019) and neuroblastoma (NB) (Kildisiute *et al*, 2021). We ran the algorithm for each data set individually and merged the resulting regulatory tables (Figure 4A, Figure S2, Supplementary Table 2). In the merged regulatory table (Figure 4A), the rows correspond to regulators (TFs) and the columns to gene modules derived from each individual study. The top annotation bars indicate the disease type and study that each module originate from. It is evident that certain meta-modules emerge (Box 1), consisting of several modules with similar gene content from different studies that are regulated by the same regulators.

**Figure 4.**
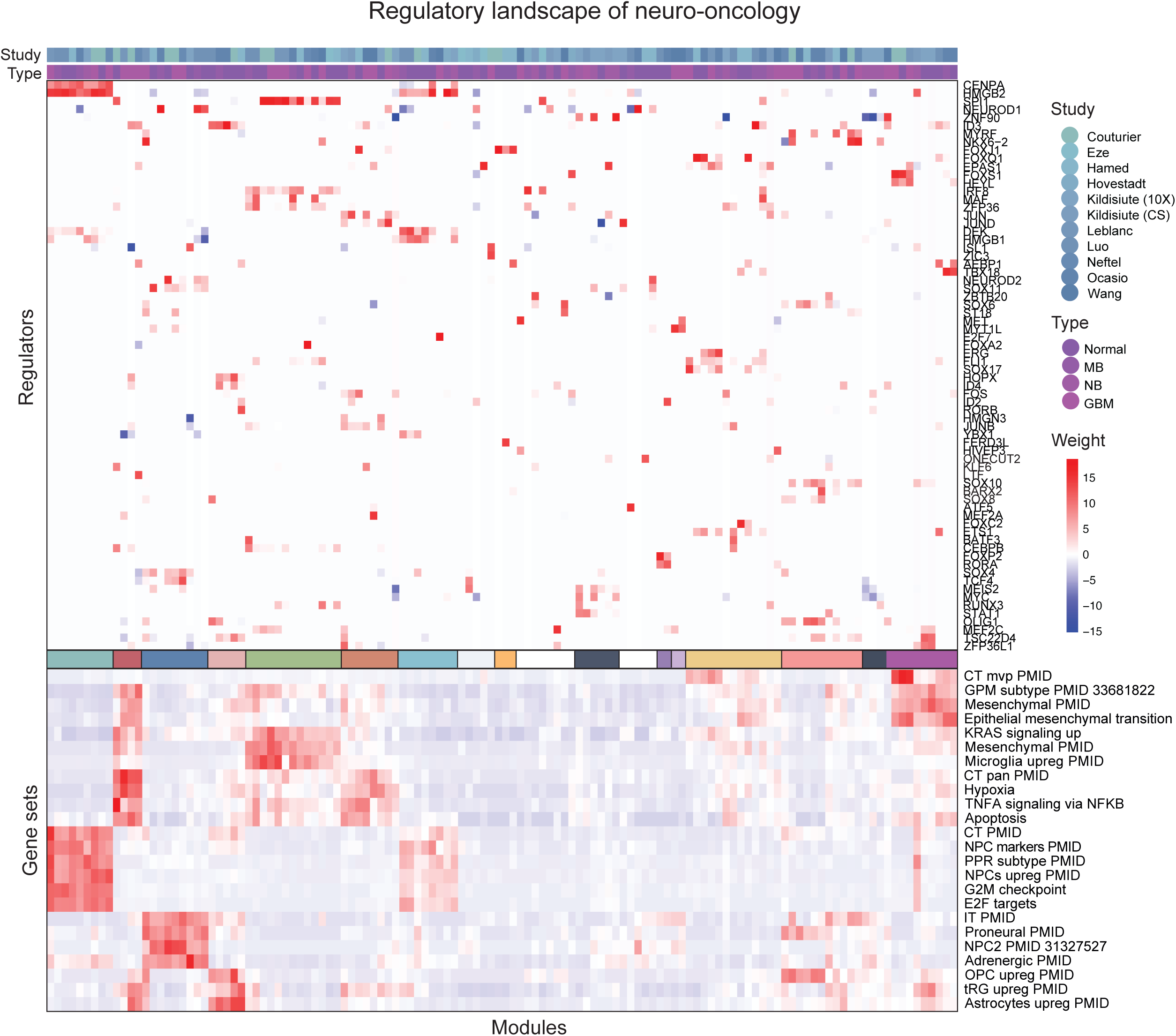
The regulatory landscape of neuro-oncology. Middle panel is the regulatory table from *scRegClust*, with modules as columns and regulators (TFs) as rows. Top panel are annotation bars indicating what type and study each module originate from. Meta-modules (Box 1) are indicated by color in the bar below the middle panel. Bottom panel display enrichments for each module against a database of neuro-oncology related gene sets, derived from studies indicated by PubMed ID (PMID). Abbreviations: CT = cellular tumor, MVP = microvascular proliferation, GPM = glycolytic/plurimetabolic, pan = pseudopalisading cells around necrosis, NPC = neural progenitor cells, PPR = proliferating progenitor cells, IT = infiltrative tumor, OPC = oligodendrocyte progenitor cells, tRG = truncated radial glia.

#### Box 1. The regulatory meta-modules.

Description of the meta-modules defined in Figure 4. The colors to the left correspond to the middle color bar in Figure 4. The regulators listed regulate at least two of the individual modules included in the meta-module and all diseases represented in the meta-module are indicated by a filled box.

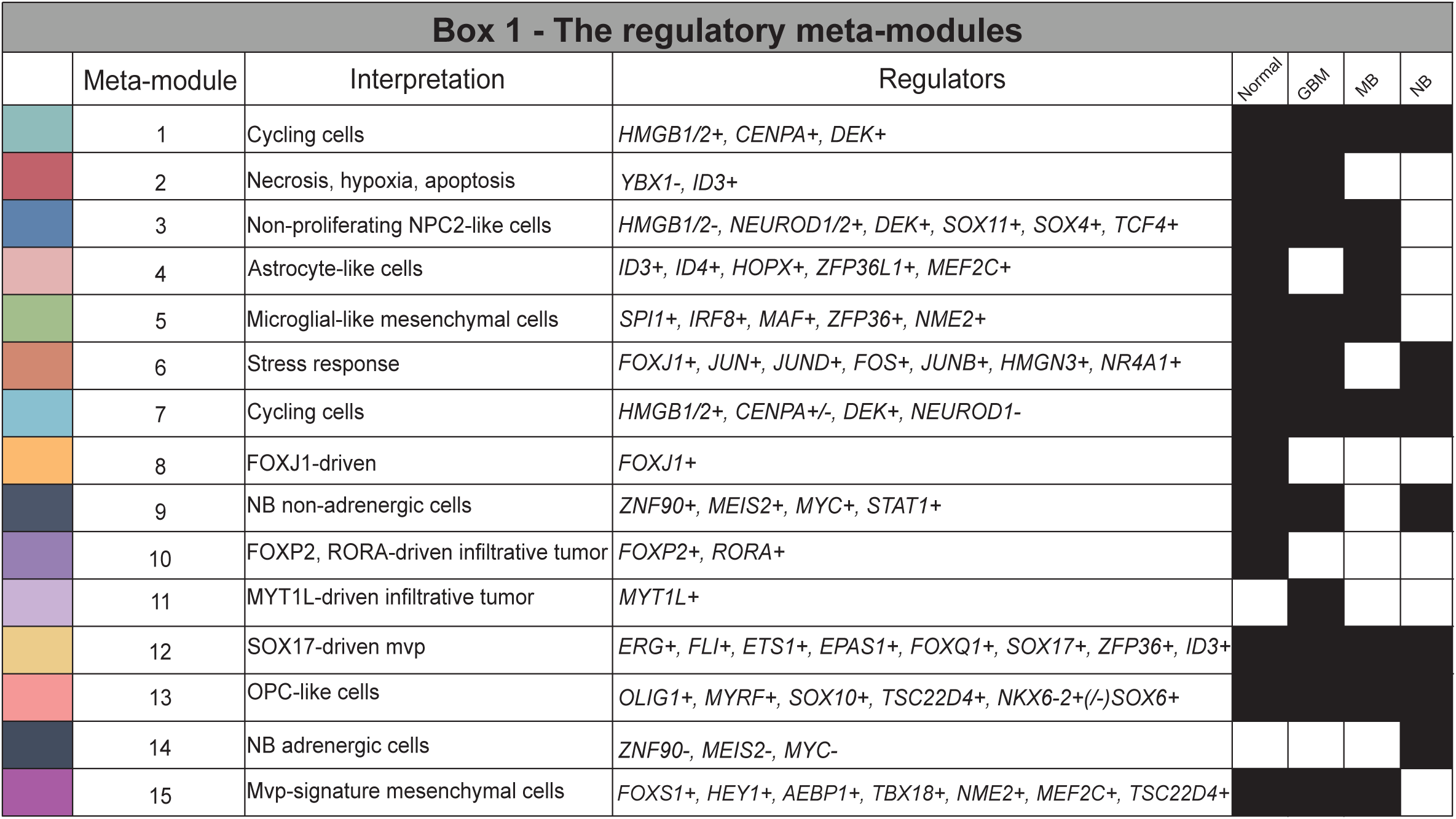

Two meta-modules, 1 and 7, represented actively proliferating cells from all types (normal tissue, GBM, MB and NB). Meta-module 1 was primarily driven by positive regulation by *CENPA* and *HMGB2*, while meta-module 7 was more strongly regulated by *HMGB1* and *DEK*. Our model suggests a subdivision of MB and normal samples along a gradient regulated by *HMGB1/2* and *DEK* vs *NEUROD1*, where *NEUROD1* promotes non-proliferating neuronal-like NPC2-like cells (meta-module 3), and *HMGB1/2* and *DEK* promotes neural progenitors with active proliferation.

Two meta-modules, 2 and 6, were enriched for hallmark signatures of hypoxia and apoptosis, as well as the Ivy atlas signature pseudopalisading cells around necrosis (CT pan), probably representing a general stress response in the cells. These were regulated by *YBX1* (#2) and members of the Fos and Jun family of TFs (#6). An astrocyte-like meta-module (#4) in normal and MB cells was linked to *ID3*, *ID4* and *HOPX*. In GBM, these cells were clustered in meta-module 2 due to their additional enrichment for stress response signatures. An OPC-like meta-module (#13) was identified that was driven by e.g. *OLIG1*, *MYRF* and *SOX10*.

There were two distinct mesenchymal meta-modules, 5 and 15. One mesenchymal meta-module (#5) coincided with an enrichment of microglial signatures and was primarily driven by *SPI1* and *IRF8* in normal, GBM and MB cells. An explanation to this meta-module is that it captures the phenotype of cells undergoing a mesenchymal shift induced by tumor-associated macrophages (TAMs) or microglia, and that the enrichment of microglia-related gene signatures in disease modules are due to this previous interaction between tumor and immune cells. *SPI1* and *IRF8* could therefore be candidate regulators of the observed mesenchymal shift in GBM cells. *SPI1* is a pioneer TF, i.e. one that can bind directly to condensed chromatin and thereafter recruit other, non-pioneer TFs, to the site. The known normal function of *SPI1* is in controlling the cell fate decision of hematopoietic cells, e.g. in macrophage differentiation. After binding to regulatory elements in the genome, it regulates gene expression by recruiting other transcription factors, such as interferon regulatory factors, e.g. *IRF8* (Pham *et al*, 2013; Le Coz *et al*, 2021). *IRF8* has been previously identified as a candidate gene involved in the immune evasion of GBM cells. Gangoso *et al* (2021) observed that *IRF8* was upregulated in GSC cells in an immuno-competent mouse model following immune attack, and since *IRF8* is normally a myeolid-specific master transcription factor, they termed this immune evasion strategy “myeloid-mimicry”.

The other mesenchymal metamodule (#15) showed no enrichment of microglial signatures, but instead showed enrichment for the Ivy atlas signature microvascular proliferation. This meta-module was interesting in the sense that it had regulators that depended on disease category. GBM samples were regulated by *FOXS1* and *HEY1*, while MB samples were regulated by *AEBP1* and *TBX18*. Normal tissue samples were driven by previously mentioned regulators but also *MEF2C* and *TSC22D4*. A third meta-module (#12) showed a slightly weaker enrichment for the microvascular proliferation signature and included all types. This was driven by several TFs from the ERG family (*ERG*, *FLI*, *ETS1*) and *SOX17*, to name a few. Several of these regulator interactions are consistent with their described role in literature, e.g. *FLI1* which is a prognostic marker in astrocytoma (Hu *et al*, 2022) and is predicted to be a regulator of perivascular-like glioma cells (Hu *et al*, 2022) and the endothelial potential of myogenic progenitors (Ferdous *et al*, 2021). Similarly, the role of *SOX17* as a promotor of tumor angiogenesis has been described in mice (Yang *et al*, 2013).

Our model also suggests a subdivision of NB samples (meta-modules 9 and 14) along an adrenal- to-mesenchymal axis, driven by *ZNF90*, *MEIS2* and *MYC*, each of which was positively linked to mesenchymal-like signatures and negatively linked to ADRN-like signatures. Remaining meta-modules (#8, 10, 11) were specific to one type and driven by just one or two regulators.

Similarly to the TF regulatory landscape, the kinase map (Figure S2) revealed several likely links and made interesting predictions. For instance, along the oligodendrocytic lineage, *AATK* is linked to a meta-module in both GBM and normal developing brain that is enriched for oligodendrocyte differentiation. *PDGFRA* and *DGKB* is linked to a meta-module with an OPC-like profile, whereas *PDGFRB* is linked to invasiveness and microvascular profileration signatures. *CKB* regulates a meta-module of NPC signature targets in normal developing brain, GBM and MB, and suppresses cell cycle genes in MB and GBM. *CCND1* suppresses NPC markers. Mature astrocytes were primarily linked to *NTRK2* activity, in agreement with results in Figure 3C. *CCL2* drives a microglia-like signature in both normal developing brain and MB. As in the TF map, known cell cycle driving kinases converge on similar modules in GBM, MB and normal developing brain, e.g. *AURKA*, *PLK1* and *CCNB2*. Still there was an MB-specific *TK1* driven cell cycle module. *AK4* links to a GBM-specific module enriched for cellular tumor genes. We also found some hits of unclear significance, like *SGK1* regulating a common meta-module in GBM, MB and normal developing brain.

## Discussion

The high extent of intratumoral heterogeneity in nervous system cancers and the plastic behavior of tumor cells are major obstacles in the search for more efficient therapies against these often deadly diseases. The enormous amount of scRNA-seq data that has been generated to map intratumoral heterogeneity presents great opportunities to understand the regulation of transcriptional cell states and cell state transitions, but it also challenges us to develop suitable methods to properly analyze the data. In this work, we have addressed the lack of computational methods to identify such regulators of intratumoral heterogeneity and present a method that simultaneously detects gene modules and their regulators from scRNA-seq data. The utility of our method is demonstrated through three use cases, where it is applied to real data, and through a thorough investigation of model properties using synthetic data.

By associating a regulator model with each module, *scRegClust* improves substantially on initial clustering results derived from the correlation of target genes and regulators. We further developed scores, such as R^2^ per module, regulator importance, and a silhouette score, to help the user with parameter choices and to support the interpretation of the resulting regulatory network. In our first use case, we apply *scRegClust* to the PBMC data set and compare our results to those of SCENIC+. We show that we can, in an unsupervised fashion, recapitulate signatures of the known cell types present in the data as well as several known regulators of each cell type. The result is comparable to that of SCENIC+, but obtained in shorter time and with only one data source. We do however emphasize that although the methods produce a similar output, they differ in several important technical aspects, and should be treated as complementing rather than competing methods.

In our second and third use case, *scRegClust* is applied to data sets from nervous system cancers (GBM, MB and NB) and the developing brain and used to derive insights about the diseases that warrant further investigation. We identify several regulators of an OPC-like, invasive GBM state and combine temozolomide and dasatinib, a tyrosine kinase inhibitor targeting the state regulators, to potentiate TMZ treatment. Dasatinib has in previous studies shown anti-migratory effects (Li *et al*, 2015), consistent with state 5 having an invasive profile, and were tested in combination with temozolomide in a clinical trial as treatment against glioblastoma (Milano *et al*, 2009). However, this trial was terminated before proceeding to phase II. We find that the combination treatment is only synergistic in our U3065MG cell line when we first pre-treat the cells with dasatinib, followed by combined treatment with both drugs, and speculate whether this means that state 5 first needs to be depleted for the combination treatment to be synergistic. These speculations need to be more carefully investigated by measuring whether the protein levels of state 5 markers decrease following dasatinib treatment. Second, by performing an integrative study of the regulatory programs of GBM, MB, NB and the developing brain, we find that *SPI1* and *IRF8* are candidate regulators of the mesenchymal shift induced by TAMs and/or microglia in GBM. These regulators can be an important piece of the puzzle to understand the mechanism behind the observed mesenchymal transition, which appears to be a strategy for the tumor cells to evade the immune system and become increasingly resistant to treatment.

For future developments, we see several possible extensions of the algorithm. At present, *scRegClust* has the functionality to include prior knowledge of gene-gene relationships to guide the clustering of target genes. It would however be interesting to explore the possibility of including prior knowledge in the regulator-target gene relationship as well, e.g. by favouring connections that have support in an external data set (ATAC-seq, CHIP-seq etc). In addition to this, we see extensions related to non-additive contributions to the regulation of a module, extended regulation modeling to allow for feedback regulation, or automatic partitioning in regulator- and target genes.

To conclude, we present an algorithm that operates on scRNA-seq data alone to construct regulatory programs consisting of regulators and target gene modules, and use it to understand the regulatory mechanisms of cell state plasticity. The algorithm is provided as an easy-to-use R-package and can be applied to any scRNA-seq data set.

## Methods

### Datasets

All publicly available data sets used in this publication are listed in Table 1, including the data repository from where they were downloaded. The notation “3CA” refers to the Curated Cancer Cell Atlas provided by the Tirosh Lab (Gavish *et al*, 2021).

**Table 1:** Table of data sets used in this publication.

| Study | Data repository | Disease |
| --- | --- | --- |
| Couturier et al. 2020 | 3CA/brain | Developing brain |
| Couturier et al. 2020 | 3CA/brain | Glioblastoma |
| Darmanis et al. 2017 | 3CA/brain | Glioblastoma |
| Eze et al. 2021 | UCSC Cell Browser | Developing brain |
| Hamed et al. 2022 | GEO: GSE200202 | Developing brain |
| Hovestadt et al. 2019 | 3CA/brain | Medulloblastoma |
| Kildisiute et al. 2021 | Neuroblastoma Cell Atlas | Neuroblastoma |
| Kildisiute et al. 2021 | Neuroblastoma Cell Atlas | Normal fetal adrenal gland |
| Larsson, Dalmo et al | ArrayExpress E-MTAB-9296 | Glioblastoma |
| Leblanc et al. 2022 | GEO: GSE173278, GSE193884 | Glioblastoma |
| Luo et al. 2021 | Short Read Archive: GSE156633 | Developing cerebellar granule |
| Luo et al. 2021 | Short Read Archive: GSE156633 | Medulloblastoma |
| Manno et al. 2021 | Sequence Read Archive: PRJNA637987 | Developing brain |
| Neftel et al. 2019 | 3CA/brain | Glioblastoma |
| Ocasio et al. 2019 | GEO: GSE129730 | Medulloblastoma |
| Ocasio et al. 2019 | GEO: GSE129730 | Cerebellum |
| Wang et al. 2020 | 3CA/brain | Glioblastoma |
| Weng et al. 2019 | GEO: GSE122871 | Developing brain |
| PBMC | 10X website | human blood cells |

### Data processing and formatting

If available, raw counts were downloaded from the indicated source in Table 1. The data was processed before further analysis in R (R Core Team, 2022) using the package Seurat (Butler *et al*, 2018). Cells containing less than 200 genes and genes present in less than 3 cells were filtered out. Data generated from a UMI-based single-cell sequencing protocol (10X Chromium, CEL-seq2) were normalized using sctransform (Hafemeister and Satija, 2019), while expression levels for remaining data sets were quantified using log_2_(T PM/10 + 1), as suggested by Tirosh *et al* (2016). When needed, non-malignant cells were filtered out from the data set based on the provided metadata from the authors or the 3CA-portal (Gavish *et al*, 2021). We provide functionality to format the normalized data matrix to conform with the desired input of our *scRegClust* -algorithm (more details below).

### Clustering Algorithm

We developed a two-step alternating algorithm to simultaneously perform clustering on a set of target genes as well as to associate a set of regulators with each cluster (module). In a first step, given an initial clustering of the data, the algorithm determines the linearly most predictive regulators for each cluster as well as whether the regulator acts stimulating or repressing. Then, in a second step, these optimal regulators are used to allocate target genes into clusters. Prior knowledge about target gene relationships can be included to guide cluster allocation. These two steps are repeated until configurations stabilize or a maximum number of steps is reached. The algorithm was implemented as an R-package (R Core Team, 2022) and can be found on GitHub at https://github.com/sven-nelander/scregclust.

#### Algorithmic outline of *scRegClust*

The algorithm takes as its input two matrices **Z***_t_* and **Z***_r_* containing *n* rows of matching observations, typically cells, on *p_t_* target genes and *p_r_* tentative regulator genes in their columns, respectively. In addition, the algorithm requires the number *K* of desired clusters and an initial clustering of the target genes into *K* clusters. If the latter is not supplied, an initial clustering is produced as described below under “Preprocessing and initialization”.

Formally, our approach aims to solve the clustering problem by determining

1. a *K* × *p_t_* matrix of cluster membership **Π**, such that **Π**^(*i,j*)^ = 1 if target gene *j* is in cluster *i*, and 0 otherwise, (equivalently, let C*_i_* be the set of all target genes in cluster *i*)
2. a subset R*_i_* of regulators associated with each cluster, where |R*_i_*| ≤ *N_r_*,
3. a vector of signs ***s****_i_* of size |R*_i_*| for each regulator associated with a cluster,
4. non-negative coefficients **B***_i_* of dimension |R*_i_*| × *p_t_* for each cluster,
5. positive variance parameters 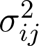 for each gene and cluster, such that the following target function is minimized

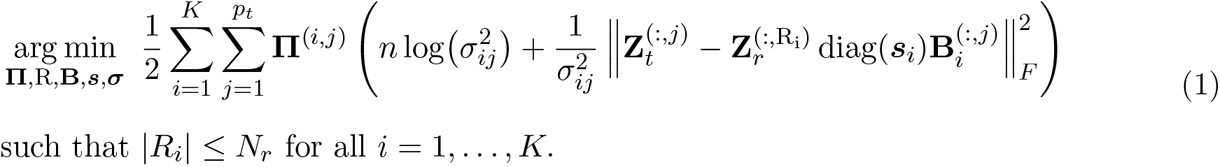

Note, that we do allow regulators to appear in multiple clusters. Additionally, we make the assumption that a regulator acts stimulating (positive sign) or repressing (negative sign) on all target genes in the cluster.

The direct solution of this optimization problem, however, requires to check a combinatorial number of regression problems, which is infeasible in practice. To alleviate this, we instead used a data-driven approximation to solve Eq. (1).

##### Algorithm 1 High-level overview of the *scRegClust* algorithm.

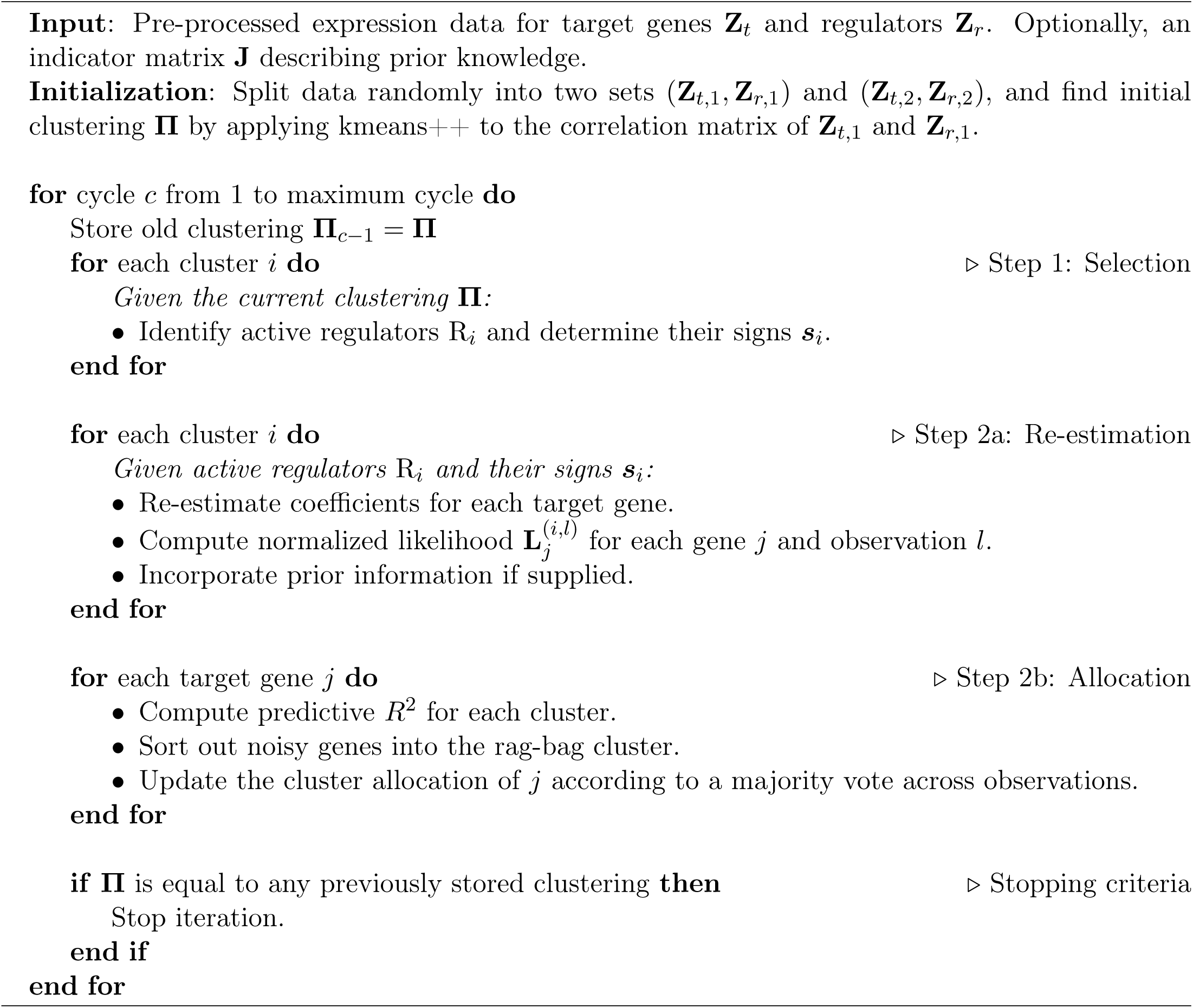

#### Data-driven approximation

To make the computation possible, we proceed in the fashion of alternating clustering algorithms, such as k-means, by alternating between (1) determining the structure of clusters, here determining R*_i_*, ***s****_i_*, and **B***_i_* for each *i* as well as 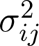 for each *i* and *j*, and (2) re-assigning cluster membership, here determining **Π**.

##### Preprocessing and initialization

Observations of target genes and regulators (or a randomly reduced subset) are randomly split into two sets to allow for unbiased estimation of the model quality during cluster allocation below. The ratio is user-defined but 50-50 by default. If the observations are stratified, the sample assignment of each observation can be supplied and splitting is performed within each sub-group. We write **Z***_t,i_* and **Z***_r,i_* for the i-th data split containing *n_i_* observations (*i* = 1, 2). The first data split is regarded as a training set, whereas the second data split functions as a test set. We therefore compute the preprocessing below on the training set and apply the same values to the test set.

During data splitting, the user can choose whether or not to center the target genes within each sub-group defined by the sample stratification. In addition, after splitting, both target and regulatory genes are centered individually (without regard to stratification) and regulatory genes are scaled to standard deviation 1.

If no initial cluster membership is provided, the cross-correlation matrix of **Z***_t,_*_1_ and **Z***_r,_*_1_ is computed. The k-means++ algorithm (Arthur and Vassilvitskii, 2007) with the cross-correlation matrix as an input and multiple restarts is then used to find an initial clustering of the target genes into K clusters. Note that this step already takes regulators into account and is therefore not equivalent to the clustering of target genes.

##### Determining a regulatory model for each cluster

Given the cluster membership of target genes, the goal of Step 1 is to determine which regulator are linearly most predictive for the target genes in each cluster. Since cluster membership is considered known, the optimization problem in Eq. (1) can be solved separately for each cluster. However, due to the computational complexity of the optimization problem, it is not possible to solve it exactly. In our scenario, there are 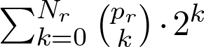candidate sets of regulators and their signs to consider. Instead, we used the cooperative-Lasso (Chiquet *et al*, 2012, coop-Lasso) to find an approximate solution to the problem. The coop-Lasso solves the following optimization problem

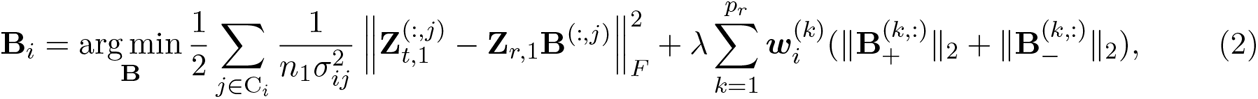

where λ is a penalty parameter related to *N_r_* above, controlling the amount of regulators that will be selected; ***w****_i_*is a vector of weights that can be cluster-specific and will be described below; **B**^(*k,*:)^ refers to the k-th row in **B**; 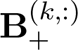 and 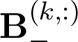 are defined by setting all negative or all positive elements in **B**^(*k,*:)^ to zero, respectively.

The coop-Lasso selects which regulators, or technically, which rows of **B**, are included in the model by setting the coefficients of the deselected groups to zero, similar to the group Lasso (Yuan and Lin, 2006). In addition, the coop-Lasso aims for sign-coherence in each group and induces sparsity within groups, deselecting target genes not affected by some of the regulators. This means that typically the rows of the estimated coefficient matrix **B***_i_* will have all positive or all negative sign. However, if coefficients are close to zero or *λ* is small it is possible that coefficients within one group have mixed sign. To determine the sign of each regulator, we therefore assign 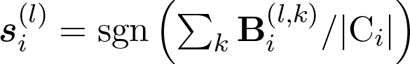 set of active regulators R*_i_* is equal to those corresponding to non-zero rows of **B***_i_*. This way, the coop-Lasso in Eq. (2) provides an approximation to the optimization problem in Eq. (1) for fixed cluster membership.

To make computations on a matrix of coefficients efficient, we apply over-relaxed Alternating Direction Method of Multipliers (ADMM) (Boyd *et al*, 2011) to the coop-Lasso problem, splitting the variables such that one set is specific to the loss and the other is specific to the penalty, leading to simple solutions for each separate sub-problem. To solve the sub-problem corresponding to the penalty, we use an explicit form of the proximal operator of the coop-Lasso given in Chiquet *et al* (2012). To speed-up convergence we compute ADMM step-length and over-relaxation parameters in an adaptive fashion as described in Xu *et al* (2017).

Only the coefficients **B***_i_* are optimized in Eq. (2). The variance parameters 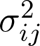 are treated as known and weights are assumed to be given. We use a plug-in estimate for the variance parameters computed using an ordinary least squares (OLS) estimate 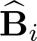 of the regression coefficients of 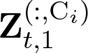 on **Z***_r,_*_1_. They are estimated in an unbiased way as

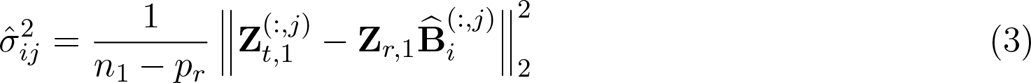

To debias the estimated coefficients, weights are selected in a fashion similar to the adaptive Lasso (Zou, 2006) before estimation of **B***_i_*. Setting the weights to

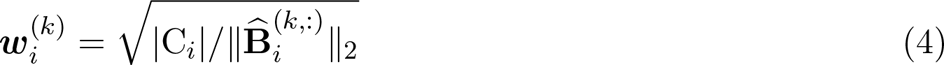

improves the selection of regulators.

##### Determining cluster membership

In Step 2, cluster membership is re-allocated, based on the updated cluster structure determined in Step 1. To do so requires the estimation of non-negative coefficients **B***_i_* for each cluster, estimation of residual variances 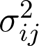 for each gene and cluster, and determination of the cluster membership matrix **Π**.

Given R*_i_*and ***s****_i_*, we used sign-constrained linear regression using non-negative least squares (NNLS) (Meinshausen, 2013) to determine the coefficients **B***_i_* for each cluster. To do so, the following optimization problem was solved

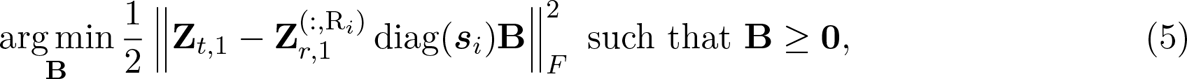

where the inequality is considered element-wise. To compute the NNLS coefficients efficiently, we implemented the algorithm described in Nguyen and Ho (2017). We modified the algorithm to be able to perform computations on a matrix of responses instead of on a single vector. To avoid unnecessary computation, responses are excluded from the computations once they reach the desired convergence criterion. The variance parameters 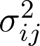 are estimated as in Eq. (3) but with the OLS estimate 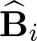 for the coefficients replaced by the NNLS estimate **B***_i_* determined in Eq. (5) and with p*_r_* replaced by |R*_i_*|.

Re-estimation of the coefficients for the selected regulators and their signs removes the bias the coop lasso introduced and enforces the chosen signs. In order to construct an allocation rule, it is necessary to estimate the coefficients for target genes that were not part of the cluster before. In the following, computations were performed on the second data split **Z***_t,_*_2_ and **Z***_r,_*_2_ to avoid bias towards the most complex regulatory programs.

Rag bag clustering is used to identify target genes that do not fit well in any cluster. To do so, the predictive *R*^2^-value is computed for each target gene and cluster from the residuals of predicting **Z***_t,_*_2_ from **Z***_r,_*_2_ diag(***s****_i_*)**B***_i_*. The best predictive R^2^ value across clusters for each target gene is recorded.

If this best value is below a user-specified threshold, then the gene is considered noise and badly predicted within all clusters. It is then placed in a noise cluster/rag bag. Only the remaining target genes are considered in the following steps.

To include prior knowledge of target genes that have a biological relationship, the algorithm allows the user to supply an indicator matrix **J** of size *q* × *q* such that **J**^(*i,j*)^ = 1 if genes *i* and *j* have a biological relationship, and zero otherwise. It is assumed that **J**^(*i,i*)^ = 0 to simplify computations below. By providing gene symbols for the target genes in **Z***_t_* and the genes in **J**, the algorithm determines the genes for which prior information is available. It is not required that the provided sets are equal as long as there is overlap. Assume for sake of notation that prior information is provided for all target genes in **Z***_t_*.

For a fixed target gene *j*, **J**^(^*^j,^*^:)^ contains 1’s for those target genes which have a biological relationship with gene *j*. Given the current cluster membership matrix **Π**, compute fractions 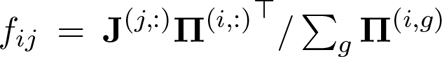 to encode the biological evidence supporting gene *j* to be in cluster *i*. In case cluster *i* is empty, set *f_ij_* = 0. These fractions are then normalized across clusters as 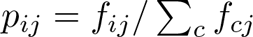. For numerical reasons, log-probabilities are used below. To avoid taking the log of zero, a small baseline parameter *α* = 10*^−^*^6^ is added to each *f_ij_* before normalization.

Given R*_i_*, ***s****_i_*, **B***_i_*, and 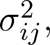, the likelihood for target gene *j* across clusters is computed for observation *l* as

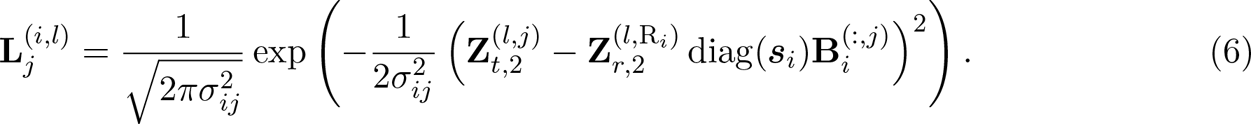

These are then normalized by setting 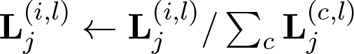. To update the cluster membership for target gene *j* we then compute votes

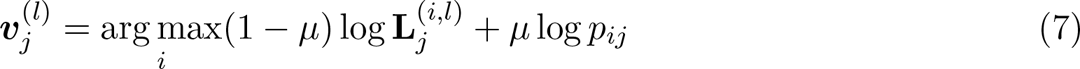

for each observation *l* and assign target gene *j* to the cluster which receives the majority of votes. The parameter *µ* ∈ [0, 1] can be used to control the strength of the prior on the allocation process.

All target genes are processed in a random ordering to avoid introducing bias. Prior fractions for each gene are computed in each iteration using the previously updated cluster assignments as well as old cluster assignments for genes that have not been updated yet.

##### Determining convergence

A history of the cluster membership matrices, **Π***_k_* for iteration *k*, is kept and the algorithm is stopped when the current **Π***_n_* computed in Step 2 is equal to **Π***_n_*_0_ for one of the previous iterations *n*_0_. At this point, the algorithm would enter a loop of length *n* − *n*_0_ and is therefore exited. We consider the algorithm as converged if *n* − *n*_0_ = 1. If *n* − *n*_0_ *>* 1, the algorithm has found an unstable cluster configuration and results for each possible configuration within the loop are returned.

#### Internal validation measures

##### Guidance on selection of the penalty parameter

To evaluate the clustering results we introduce two performance scores. The aim of our algorithm is to associate modules with their linearly most predictive regulators. As a measure of clustering quality, it is therefore natural to consider the predictive R^2^ per module, computed on the second data split with the coefficients **B***_i_* for the selected regulators re-estimated on the first data split. We therefore compute

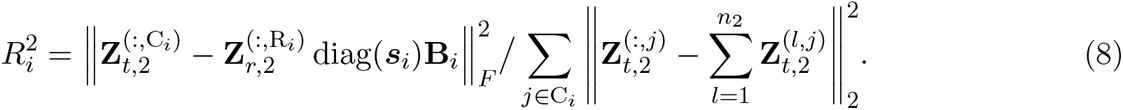

In addition, we compute the importance of each regulator within a module as follows. The regulator and sign sets without regulator *j*, denoted as R*_i,−j_* and ***s****_i,−j_*, are used to re-estimate the coefficients for module i. We then compute another 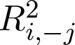 and compute the importance of regulator j in module *i* as 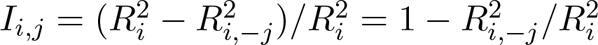. This is the ratio of the semi-partial correlation of the expression profile of regulator j with the expression profiles in module i to the overall R^2^ of module i with all regulators. A regulator that is more influential in predicting the target genes in module i will have larger importance. Importance values are typically less or equal to 1 and can be negative, which indicates that a regulator was introducing more noise than it compensated for in predictive strength.

A decrease in *R*^2^ per module with an increase in the penalization parameter *λ* is expected, due to selection of less regulators and therefore less degrees of freedom in the linear models associated with each module. Importance is the ratio of the squared marginal correlation of a regulator with the target genes in a module to the overall *R*^2^ of that module. This implies that an increase of importance values is expected to concur with a decrease in number of selected regulators. Therefore, importance is expected to increase with an increase in the penalization parameter *λ*. Selection of the penalty parameter can therefore be guided by balancing these two scores. Predictive *R*^2^ should be high while importance should neither be too low nor too high, since the latter is typically indicative of too few selected regulators.

##### Guidance on selection of the number of modules

Selection of the number of modules can be aided by cross-cluster predictive *R*^2^. For each final module’s selected regulators and signs, coefficients are re-estimated for all target genes. These are then used to compute the predictive *R*^2^ for each target gene in all available modules. Denote these values as 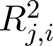 for target gene j when computed with regulators from module *i*. If there are *K* modules, this will result in a matrix of size *p_t_* × *K*. We then define the following score for each target gene *j* which has been clustered in module *c*

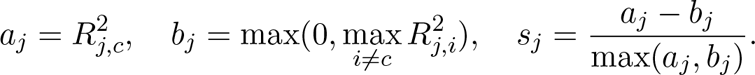

Due to the similarity with the well-known silhouette value (Rousseeuw, 1987), we call s*_j_* the silhouette score for target gene j. The average silhouette score across all target genes gives a rough measure for clustering quality and can be used as a straight-forward tool to compare different module counts. A larger average silhouette score indicates that target genes on average are better located within modules. To determine the optimal number of modules, average silhouette score should be considered jointly with predictive R^2^ per module. A good clustering achieves high values in both scores.

#### External validation measures

To evaluate the performance of simulations, we compared the groundtruth to the estimated clustering in multiple ways.

##### Adjusted Rand index

To determine overall clustering performance, we use the adjusted Rand index (Hubert and Arabie, 1985), a well-known similarity measure between two clusterings. The standard adjusted Rand index ARI does not account for rag bag clustering. We therefore consider a modified version of the adjusted Rand index

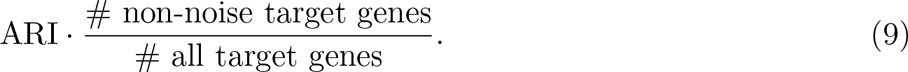

##### Regulator selection

To answer whether scRegClust selects the right regulators for each module we cannot compare regulators associated with each module directly, since estimated modules might not match clearly with the groundtruth. We therefore check correct selection of regulators for each target gene, as these are easily comparable with the groundtruth. For easier comparison we compute two measures:

1. True positive rate, as the proportion of correctly selected regulators,
2. False positive rate, as the proportion of incorrectly selected regulators.

Both of these measures are between 0 and 1 with larger values being better.

##### Implementation

The package was implemented in R (R Core Team, 2022). Computationally expensive parts were written in C++ using the packages Rcpp (Eddelbuettel and François, 2011) and RcppEigen (Bates and Eddelbuettel, 2013). The package igraph (Csardi and Nepusz, 2006) was used for visualization. The R package clusterCrit was used to compute the Rand index.

#### Drug combination treatments

##### Cell culture

The human glioblastoma cells U3065MG were obtained from the Human Glioma Cell Culture (HGCC) Biobank at Uppsala University (Xie *et al*, 2015). The cells were cultured in a mixture of Neurabasal (Gibco, #21103-049) and DMEM/F12 (Gibco, #31331-028) growth medium in 1:1 ratio, 1x B27, without Vitamin A (Gibco, #12587001), 1x N2 (Gibco, #17502001) and 1% Pen/Strep (Sigma-Aldrich, #P0781). The cells were grown as adherent culture on a laminin coated flasks (Sigma Aldrich, #L2020-1MG), and were detached for splitting and seeding by TyplE without phenol red (Gibco, #12604039). The growth medium was supplemented with 10 ng/mL recombinant human FGF-basic (Peprotech, #100-18B) and 10 ng/mL recombinant human EGF (Peprotech, #AF-100-15) each passage. The cells were regularly tested for Mycoplasma by either MycoAlert assay (Lonza, #LT07-418) or MycoStrip (InvivoGen, #rep-mys-50).

##### Cell viability assay

U3065MG cells were seeded in a laminin coated 384-well plates (Thermo Scientific, #142761) at a density of 2,000 cells/well at a volume of 0.04 mL/well. The plates were centrifuged briefly at 200 x g to collect the cells at the bottom of the plates and incubated for 24 h at 37℃ in humidified atmosphere with 5% CO2. Afterwards, the cells were either treated with temozolomide (Selleckchem, #S1237) or one of dasatinib (Selleckchem, #S1021). All compounds were dissolved in DMSO to a 10 mM stock solution. The drugs were dispensed with D300e digital dispenser (Tecan) without additional dilution of the stock solution. Afterwards, the plates were centrifuged briefly at 200 x g, and were incubated for 48 or 72 h at 37℃ in humidified atmosphere with 5% CO2. Cell viability was measured by the Alamar Blue assay (Invitrogen, #DAL1100), which was added 16 h prior the readout. After the 72 h time point, the growth medium was aspirated, and the cells were washed once with sterile PBS before addition of fresh medium. The cells were treated for additional 72 h with a combination between teamozolomide and dasatinib in a 7 × 7 dose-response matrix. 16 h prior the end point of the reaction, alamar blue was added into each well of the plates. The reduction of resazurin to resorufin was used as a proxy for determination of the cell viability. The raw measurements of the treatments were normalized to DMSO control. The BLISS coefficient and the sensitivity of the combinations were calculated with SynergyFinder (Zheng *et al*, 2022) in R (R Core Team, 2022).

## Data availability

No data was generated for this work. The R-package can be found on GitHub at https://github.com/sven-nelander/scregclust.

## Acknowledgements

We thank the Swedish Cancer Society, Swedish Childhood Cancer Foundation, Swedish Research Council and the Swedish Strategic Research Foundation for financial support.

## Author contributions

SN conceived the study. IL and FH developed the algorithm. FH wrote the R-package and IL performed the computational analyses, with support from RJ and SN. GP and AK planned and conducted the combination treatment experiments. IL and FH wrote the first version of the paper and all authors assisted in editing.

## Conflict of interest

The authors declare no conflict of interest.

## Notes

### Competing Interest Statement

The authors have declared no competing interest.

