## Appendix for "Reconstructing the regulatory programs underlying the phenotypic plasticity of neural cancers"

Larsson, Held et al. 2022

### Appendix

**Simulations** **3**

Generation of simulated data . . . . . . . . . . . . . . . . . . . . . . . . . . . . . . . . . . . . 3

Detailed simulation results . . . . . . . . . . . . . . . . . . . . . . . . . . . . . . . . . . . . . 4

**Supplementary figures** **7**

**Supplementary tables** **14**

### Simulations

#### Generation of simulated data

A suitably preprocessed scRNA-seq dataset consisting of 1912 genes and 9884 cells was modified to illustrate different aspects of real data in a controlled environment. To do so, the pairwise Pearson correlation between all genes was computed and 100 genes with maximal correlation between -0.4 and 0.4 were chosen as tentative regulators. Correlation thresholds were chosen to ensure that regulators are not collinear but retain an interesting correlation structure with similarity to real data. For each of 5 modules, between 5-15% of these regulators were then randomly chosen and associated with the respective modules. Each regulator $j$ associated with module $i$ was assigned a random positive or negative sign $\boldsymbol{s}_i^{(j)}$. To generate tentative target genes for module $i$, a mean value $\mu_{i,j}$ was chosen for each regulator $j$. Coefficients $\mathbf{B}_i$ were then simulated as described in the simulation setups below. For each module, 100 target genes were simulated. Finally, additive noise is added with signal-to-noise ratio 0.8.

**A** For each regulator $j$ in module $i$, a mean value was chosen uniformly at random between 0.01 and 0.1. Individual coefficients for each target gene were then chosen normally distributed around the chosen mean value multiplied by the regulator's sign $\boldsymbol{s}_i^{(j)}$ with standard deviation 0.1.

**B** Like **A**, however, individual coefficients were chosen with standard deviation 0.2 and if individual coefficients ended up being of the wrong sign, i.e. opposite to $\boldsymbol{s}_i^{(j)}$, they were set to zero.

**C** Like **A**, however, coefficient means were chosen between 0.1 and 0.3. As in **B**, coefficients of the wrong sign were set to zero. After simulating the true coefficients, zero-mean normally distributed noise with standard deviation 0.08 was introduced in the coefficients for all regulators, irrespective of whether a regulator was associated with a module or not.

**D** In this setup, only 3 modules were simulated with 200, 100, and 50 target genes each. In addition, 10-20% of regulators were selected per module. Otherwise, coefficients were simulated as in **A** with standard deviation 0.15.

#### Detailed simulation results

Simulation setup A tested the effect of small sign-errors, especially for small coefficients. It is reasonable to assume that regulators with small marginal effect on target genes will show up with both positive and negative correlations in the data due to chance. Therefore, we believe that this simulation setup reflects real data well. Initial clustering on the cross-correlation matrix between targets and regulators achieved an adjusted Rand index of 0.326. *scRegClust* managed to improve on this result substantially for a wide range of penalization parameters with an adjusted Rand index of up to 0.973 (Figure 2B). Figure 2A shows that predictive $R^2$ per module and regulator importance change rapidly for penalization parameters larger than 0.2. Our selection rule, presented in the article, suggests to choose penalization parameters around this change point. We leave it to the user to decide on the exact value, but generally recommend larger values to restrict the selection of regulators to more important ones. *scRegClust* showed excellent recovery of regulators in this simulated example (Figure 2B and C). In the optimal range for the penalization parameters, all selected regulators correspond to true regulators.

Simulation setup B follows the model underlying *scRegClust* closely by respecting that the coefficients for each regulator associated with a module have a unique common sign for all target genes. This led to an initial clustering on the cross-correlation matrix with adjusted Rand index of 0.675. *scRegClust* managed to improve on this result for appropriately chosen penalization parameters (Figure S3B). It is noteworthy, that the clustering result with low penalization, including all correct but also many unnecessary regulators, was essentially random (ARI 0.054 for penalization 0.004), whereas the clustering performance with the appropriate penalization can be substantially better than uninformed k-means clustering on the cross-correlation matrix (ARI 0.964 for penalization 0.2). This emphasizes the importance of choosing the penalization parameter correctly. Our selection guidelines for the penalty parameter, balancing regulator importance with predictive $R^2$ per module suggest a change point somewhere around penalization parameter 0.3 (Figure S3A). Regulator selection for appropriate penalization performed even better than for simulation setup A, which was expected due to the simulation resembling the data model more closely (Figure S3B and C).

Simulation setup C assesses the robustness of *scRegClust* in the presence of a contaminated target/regulator association structure. The initial clustering achieved an already high adjusted Rand index of 0.913, which is probably due to the increased signal strength. However, *scRegClust* managed to improve on this performance for appropriately chosen penalization (Figure S4B, ARI 0.970 for penalization 0.15). Our selection guidelines above suggest a penalization parameter between 0.075 and 0.2, when considering a region of slow and stable increase in importance without $R^2$ per module decreasing too much (Figure S4A). Since we prefer larger penalization to encourage stronger regulator selection, we recommend choosing 0.15 or 0.2 as the ideal penalization parameter. Despite increased coefficient noise, *scRegClust* uncovers most of the regulators in the groundtruth (Figure S4B). However, regulators are deselected earlier in comparison to simulation setups A and B, which can be seen as a consequence of the coefficient noise (Figure S4C).

Simulation setup D tested the effect of unequal module sizes. Clustering on the correlation matrix between target genes and regulators resulted in an adjusted Rand index of only 0.062. *scRegClust* improved substantially on this result for appropriately chosen penalization (Figure S5B, ARI 0.963 for penalization 0.1). This shows that the integrated approach of structure modelling and clustering is superior to clustering on the correlation structure. Using our guidelines for selection of the penalty led to an obvious choice around 0.15. The quality of regulator recovery improved with the number of target genes in a module (Figure S5C). For the large Module 1, most regulators not selected in the groundtruth vanished quickly for increasing penalization. Module 2 was mid-sized and showed a slight increase in false positives being deselected by the algorithm slightly slower in comparison to Module 1. For the small Module 3, regulator selection was much noisier with respect to the groundtruth and the chance of false positives increased, with one false positive remaining even in the optimal range of penalization parameters.

The impact of under- and overspecification of input parameter $K$, setting the desired number of modules, and the effect of the minimum cluster size setting in case of cluster overspecification was shown in another simulation study based on simulation setup A, which contains 5 modules. In this dataset there are 500 target genes with 100 target genes in each module. We chose to configure *scRegClust* to allocate 2, 5, and 10 modules, respectively. During runs with 10 modules, we also assessed the impact of minimum cluster size by setting it to 0, 20, 30, and 50. In addition, to assess the impact of randomization present in initialization and data splitting, we repeated the estimation 100 times for each setting.

We found that the choice of penalization parameter had an impact on the capability of *scRegClust* to find a stable clustering. For low penalization parameters (0.001 to 0.01) an unstable clustering result was found in up to 18% of all runs independently of the number of clusters (Supplementary Table 3). An increase in the minimum cluster size led to a small improvement in cluster stability

(11% vs 17% for 10 clusters with 0 vs 50 minimum cluster size and penalization 0.001). Up to 7 different final configurations were found for low penalization. For higher penalization, non-uniqueness in the final configuration only led to two competing clustering outcomes. Average predictive $R^2$ per cluster shows a stronger decrease after penalization 0.16 or 0.2 for all cluster count/minimum cluster size combinations. This is in line with the results in Figure 2 that indicated that the optimal penalization parameter should be chosen in the range of 0.16 to 0.2. The average adjusted Rand index shows a similar pattern which strengthens the value of our selection method. During investigation of individual runs in the case of cluster misspecification, a general pattern emerged where *scRegClust* combined different clusters with little to none regulator overlap into one cluster in case of cluster underspecification and tended to split up clusters into smaller clusters in case of cluster overspecification. When initialized to find 10 clusters, *scRegClust* found on average 8.9 non-empty clusters compared to 5 in the groundtruth (penalization 0.16, Supplementary Table 3). Specifying a minimum cluster size decreased the splitting of groundtruth clusters into smaller clusters. The reduction appears gradually from 8.9 to 4.9 when minimum cluster size is increased from 0 to 50 with penalization at 0.16. The number of target genes classified as noise increases with an increase in the penalization parameters (from 2 to 374 in the case of two clusters; similar but less extreme in other settings). This is to be expected, as less regulators are associated with each cluster, and regulator models become less flexible to predict the behavior of target genes.

Another tuning parameter of algorithm is the initial number of modules $K$. As shown above, under- or overspecifying this number can have substantial effects on the final clustering result. To guide the selection of this parameter, we introduced the silhouette score (see “Methods”). We used *scRegClust* to determine the clustering result for data from Simulation Setup A, when $K$ was chosen in the range 2 to 8 (Figure S6). When looking at the average predictive $R^2$ per module, it is clear that configurations with 5 and 8 final clusters achieve high scores (Figure S6 A and B). The average silhouette score decreases only slowly from $K = 2$ to 5, and decreases more rapidly afterwards. Especially, the average silhouette score for $K = 8$ is substantially lower than for $K = 5$. Combing these two scores indicates that $K = 5$ is the most appropriate choice for the number of modules.

#### Supplementary figures

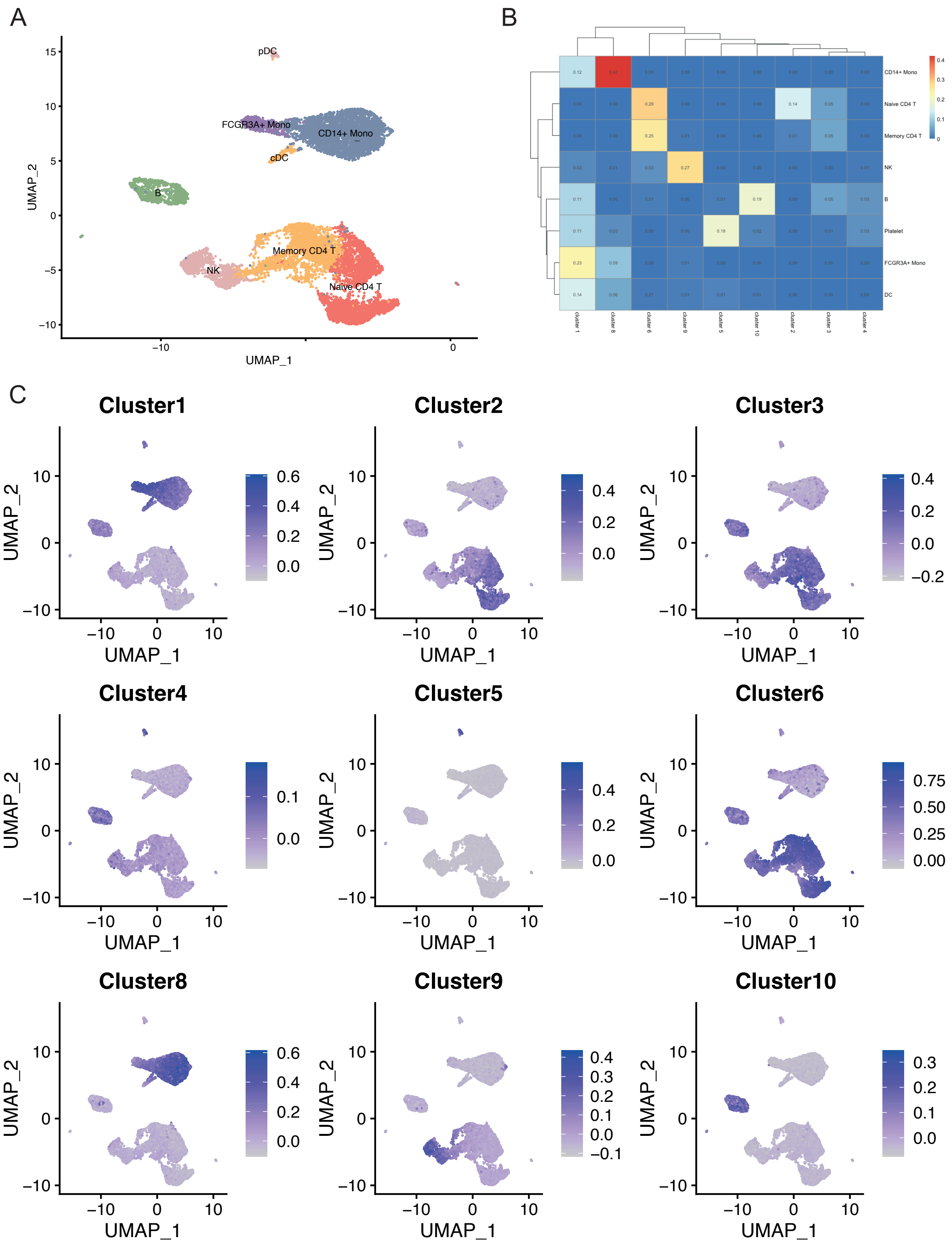


**Figure S1 Mapping of module markers to markers of peripheral blood mononuclear cell types.** A) UMAP of cell types, B) overlap between module markers and cell type markers quantified using Jaccard index, C) cells scored against module signatures and plotted using FeaturePlot() in Seurat.

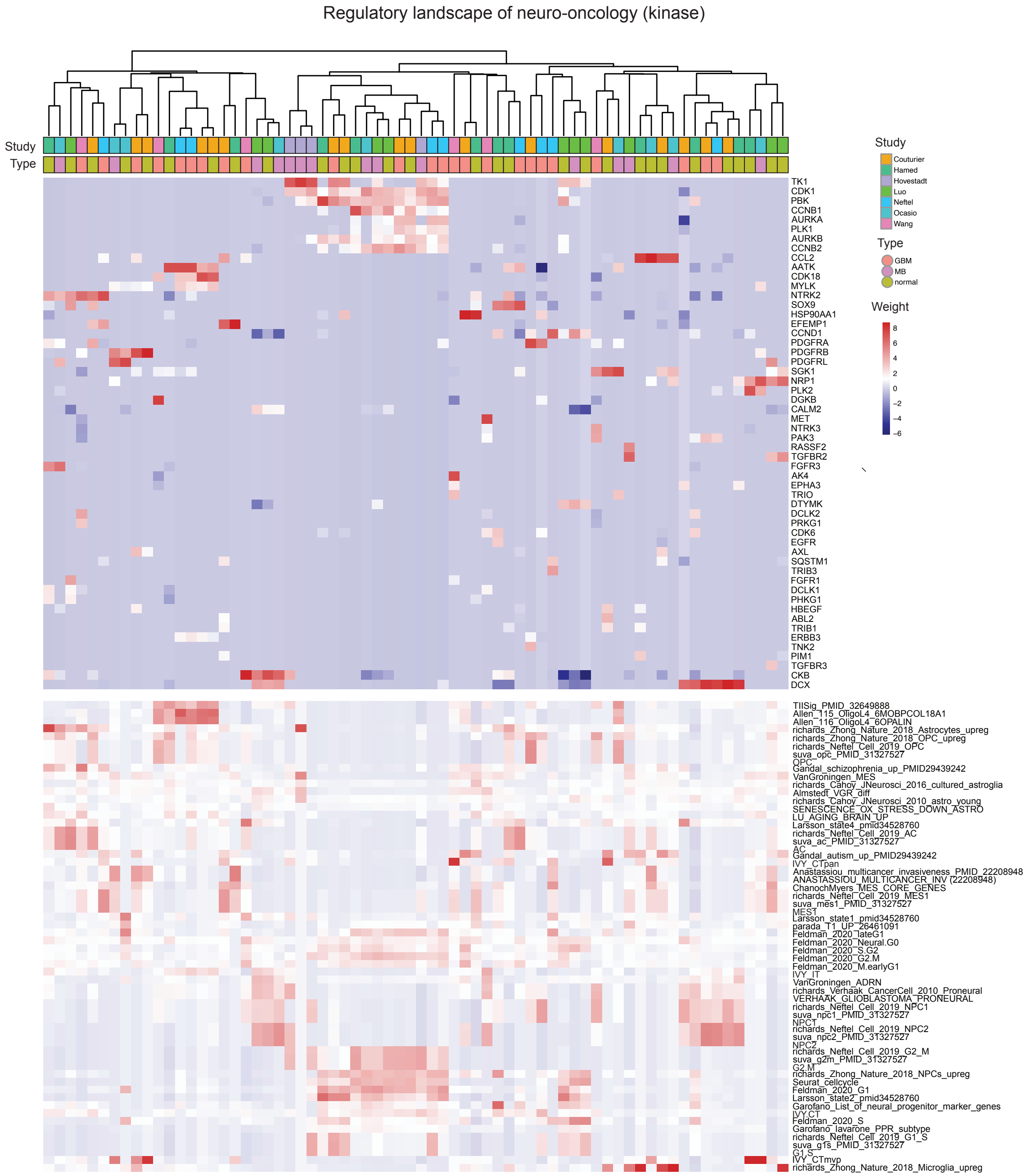


**Figure S2 The regulatory landscape of neuro-oncology - kinase version.** The figure is analogous to Figure 4A, but here the algorithm has been run in kinase mode and excluding neuroblastoma-samples. Middle panel is the regulatory table from *scRegClust*, with modules as columns and regulators (TFs) as rows. Top panel are annotation bars indicating what type and study each module originate from. Bottom panel display enrichments for each module against a database of neuro-oncology related gene sets.

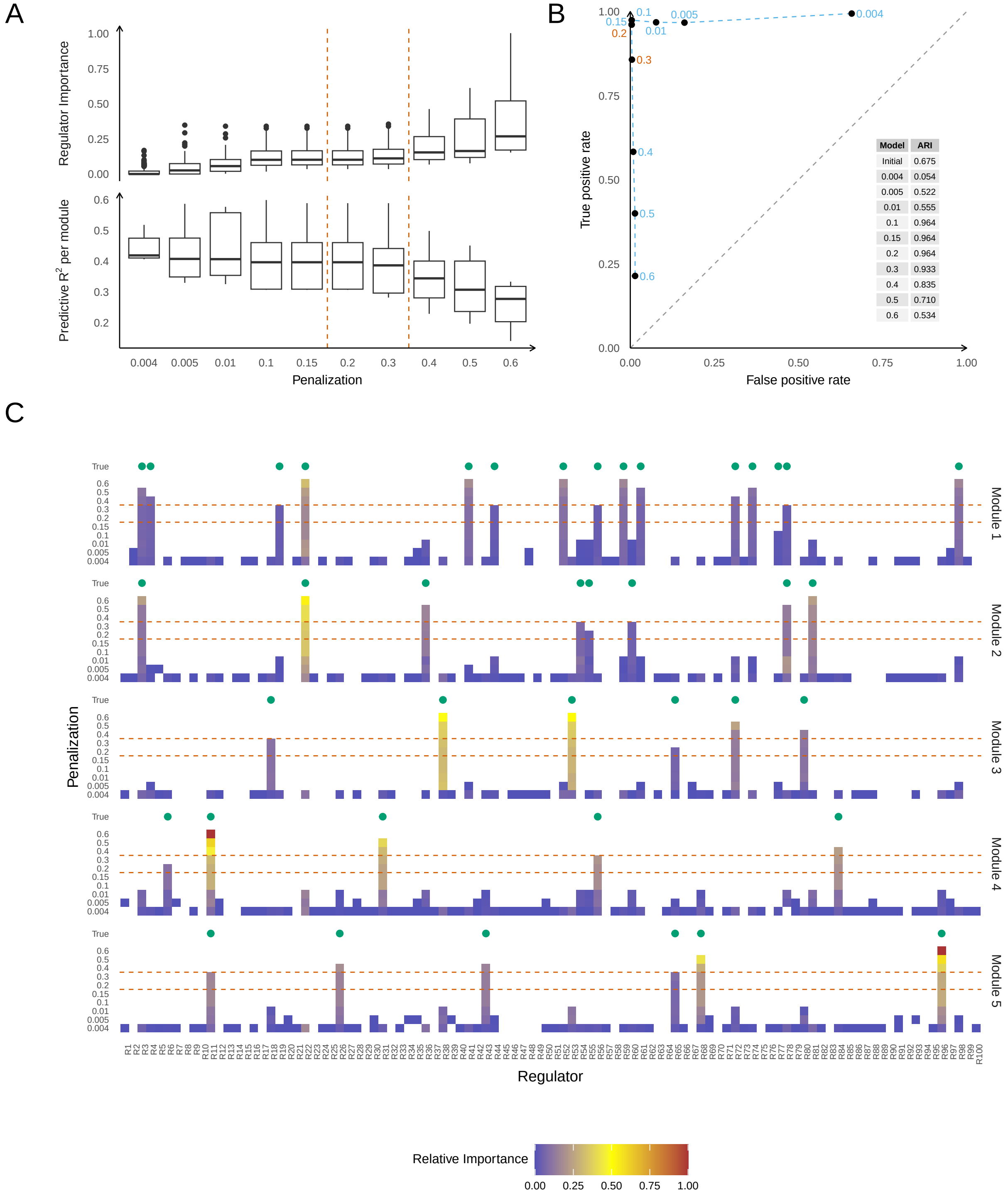


| Model | ARI |
|---|---|
| Initial | 0.675 |
| 0.004 | 0.054 |
| 0.005 | 0.522 |
| 0.01 | 0.555 |
| 0.1 | 0.964 |
| 0.15 | 0.964 |
| 0.2 | 0.964 |
| 0.3 | 0.933 |
| 0.4 | 0.835 |
| 0.5 | 0.710 |
| 0.6 | 0.534 |

**Figure S3** This figure is analogous to Figure 2 but for simulation setup B.

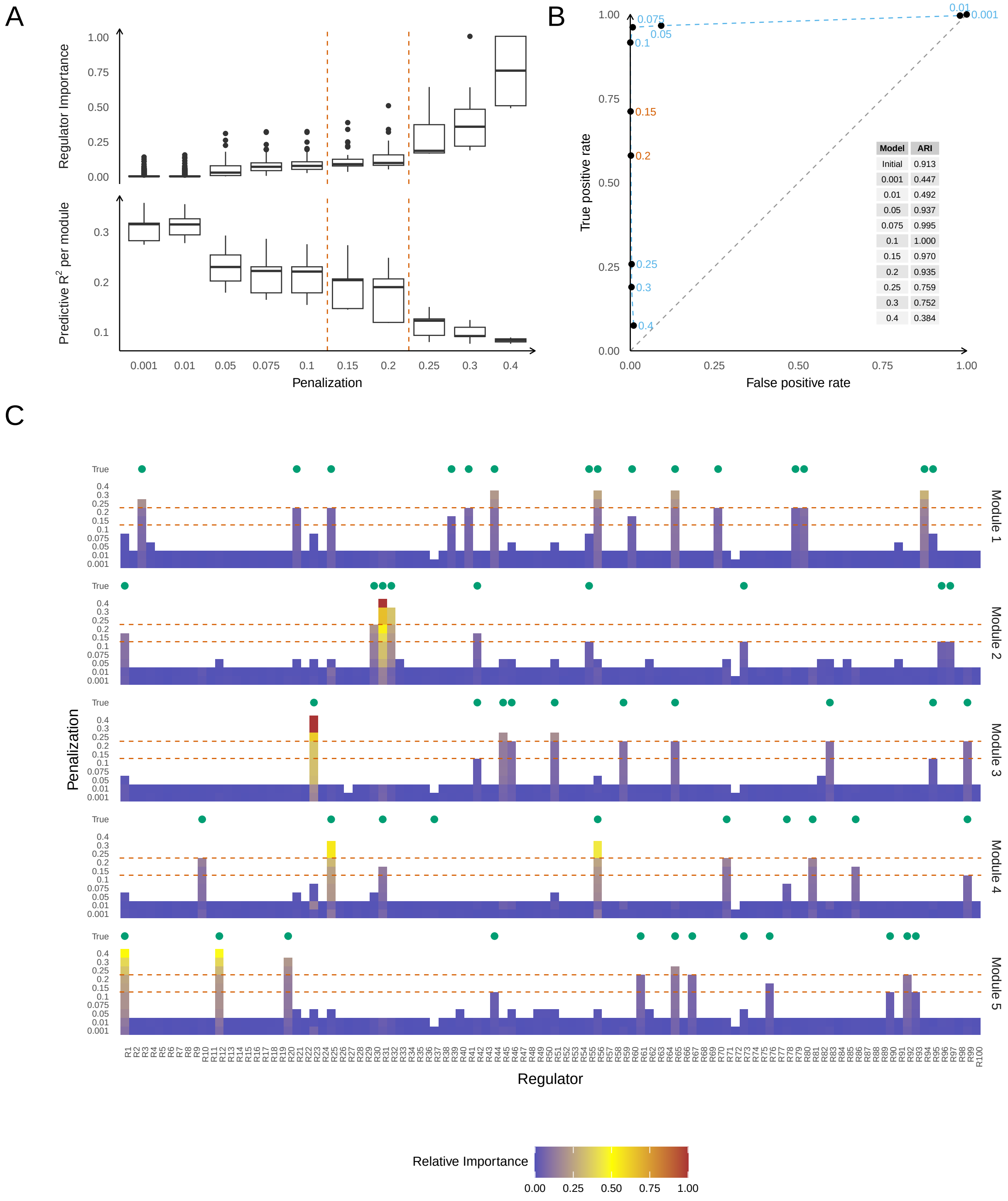


**Figure S4** This figure is analogous to Figure 2 but for simulation setup C.

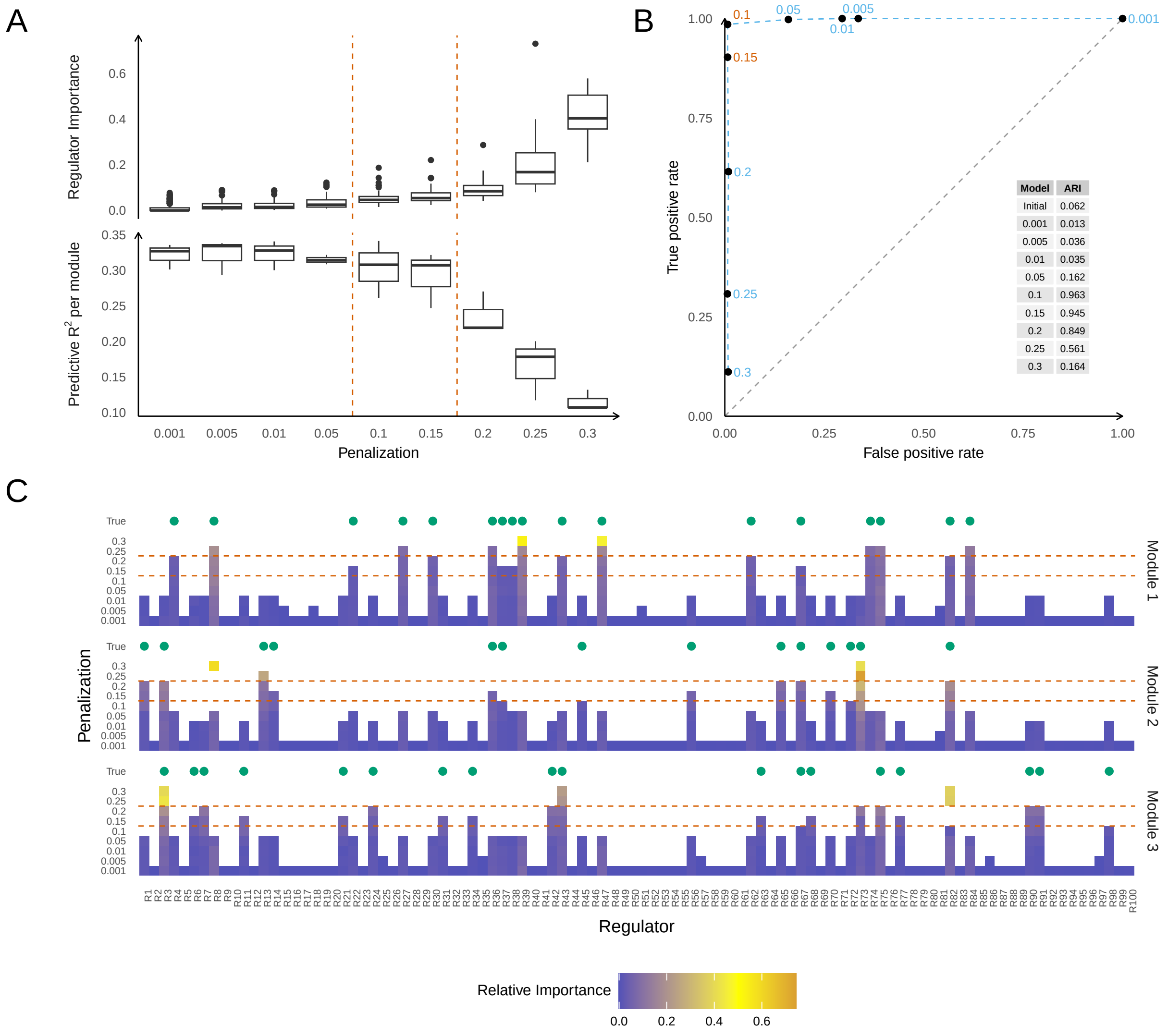


**Figure S5** This figure is analogous to Figure 2 but for simulation setup D.

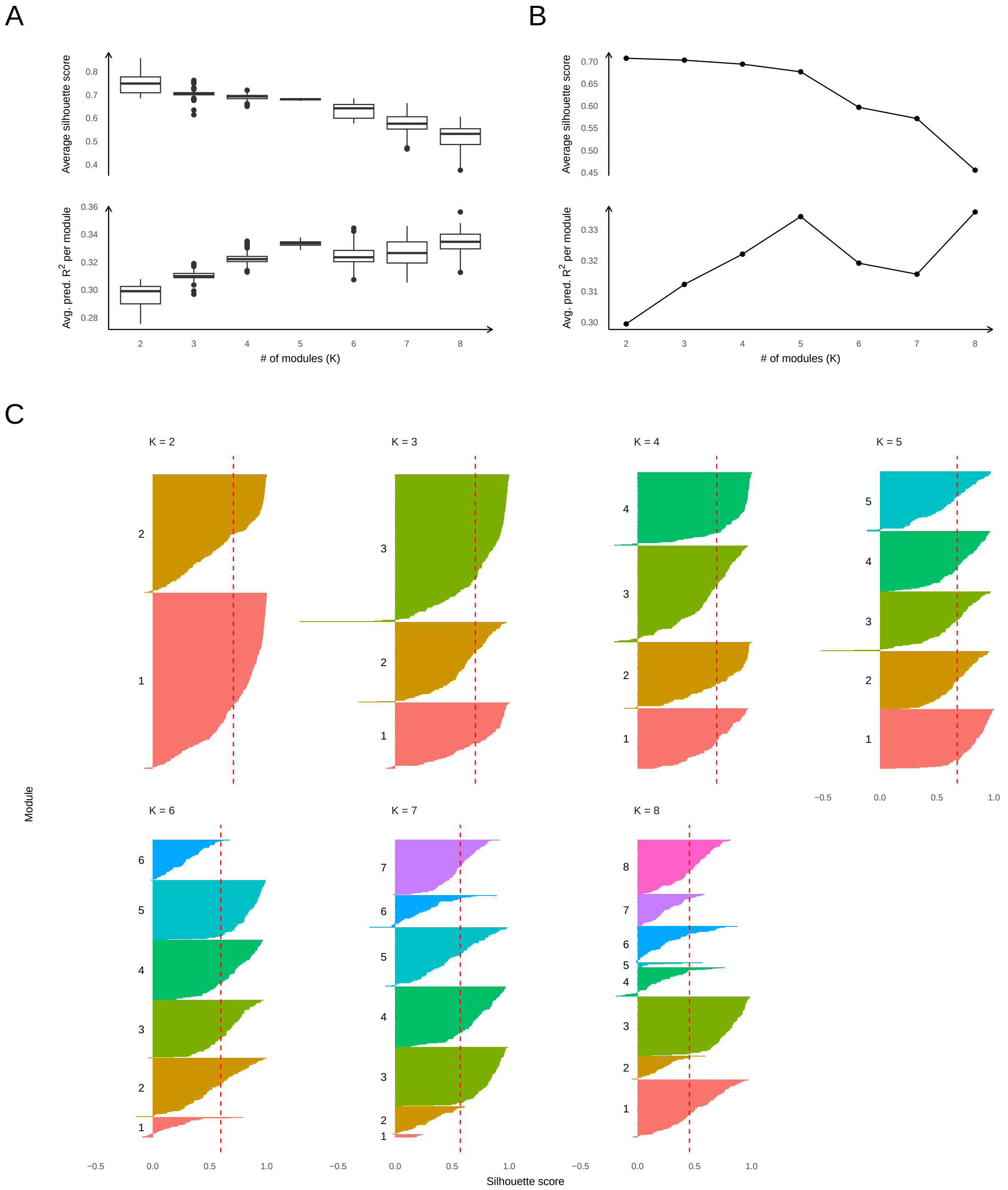


**Figure S6** Part A shows boxplots of the average silhouette score and the average predictive $R^2$ per module for a range of module counts $K$ across 100 repeated runs of *scRegClust* on data from simulation setup A for fixed penalization parameter 0.16. Part B illustrates a representative run. Part C shows the silhouette score for each module count for the run in Part B. Dashed red lines indicate the average silhouette score. Target genes have been grouped by module and sorted by decreasing silhouette score. Colors and labels to the left of each group indicate the module.

### Supplementary tables

#### Supplementary Table 1

#### Supplementary Table 2

100 runs of *scRegClust* were performed for each setting on data from simulation setup A. Compared to the groundtruth module count, which is 5, modules were underspecified (2 modules), overspecified (10 modules), and specified correctly (5 modules). In addition, minimum cluster size for 10 modules was either 0, 20, 30, or 50.
